# DJ-1 mediates regulation of metabolism and immune response in Parkinson’s disease astrocytes and Glioblastoma cells

**DOI:** 10.1101/2024.10.31.621212

**Authors:** Pauline Mencke, Jochen Ohnmacht, Félicia Jeannelle, Adrien J Ries, Mónica Miranda de la Maza, Mathilde Ullrich, François Massart, Patrycja Mulica, Katja Badanjak, Zoé Hanss, Arkadiusz Rybicki, Paul Antony, Sylvie Delcambre, Giuseppe Arena, Gérald Cruciani, Javier Jarazo, Floriane Gavotto, Christian Jäger, Anouk Ewen, Maria Pires Pacheco, Dirk Brenner, Jens Schwamborn, Thomas Sauter, Lasse Sinkkonen, Gunnar Dittmar, Ricardo Taipa, David Bouvier, Johannes Meiser, Anne Grünewald, Vincenzo Bonifati, Michael Platten, Rejko Krüger, Ibrahim Boussaad

## Abstract

An inverse correlation for the expression of Parkinson’s disease (PD)- and cancer-associated genes has been previously reported. Genes that are upregulated in cancer are frequently downregulated in PD and *vice versa*. PARK7, encoding DJ-1, was initially identified as an oncogene, but loss of DJ-1 causes early-onset PD. However, it remains elusive how differential DJ-1 levels contribute to opposite cell fates in cancer and PD. Here, we demonstrate specific effects of differential DJ-1 protein levels on the energy metabolism and cell growth in patient-derived cellular models of PD and glioblastoma (GBM) cell lines. Impaired energy metabolism was associated with an increased immune response upon IL-1β stimulation and increased apoptosis and decreased cell growth in models of PD, whereas in GBM cells increased metabolic activity translated into a reduced immune response and increased cell growth. Furthermore, we found decreased glutathione (GSH) synthesis and therefore increased levels of reactive oxygen species (ROS) and oxidized glutathione (GSSG) in models of DJ-1 deficiency and decreased ROS levels in GBM cell lines. Thus, the mechanism by which DJ-1 modulates these phenotypes is the same in both diseases.

**Graphical Abstract:** 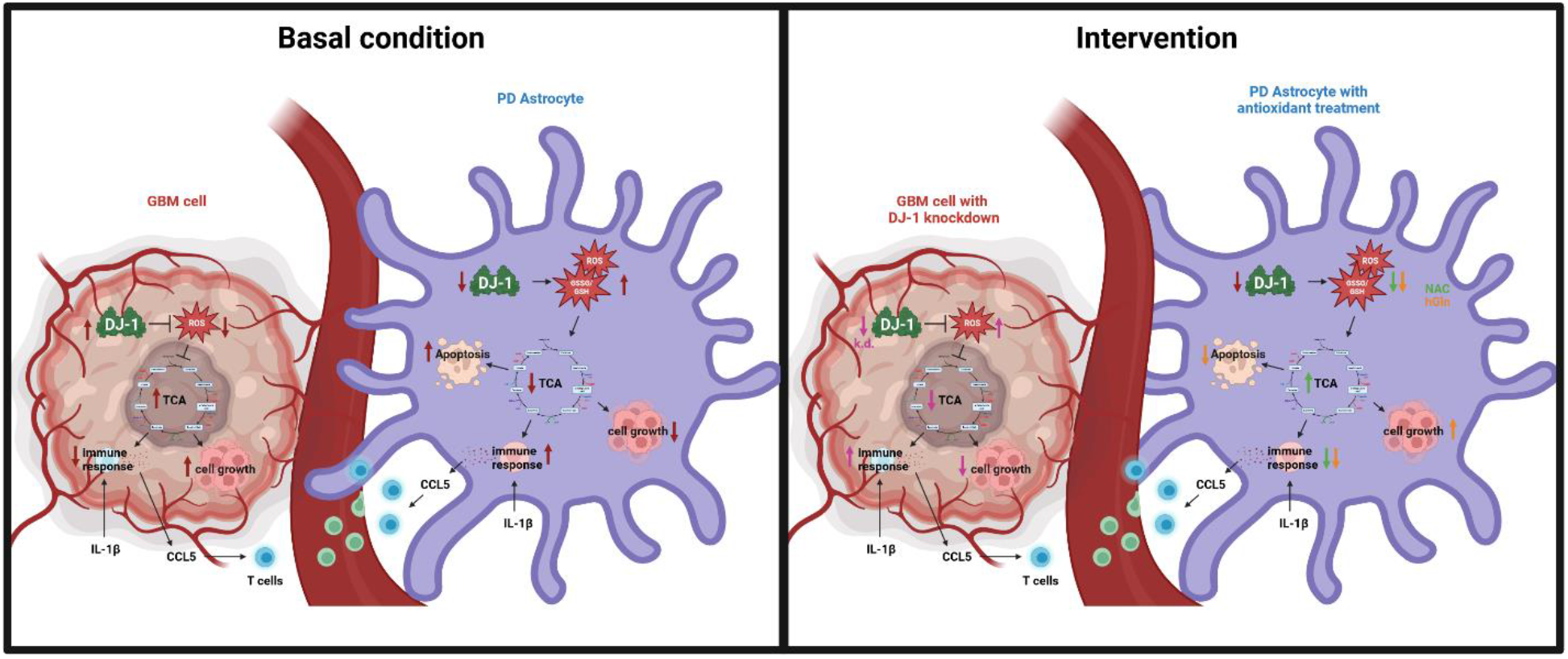

DJ-1 levels modulate GSSG/GSH ratio and ROS levels, which results in divergent effects on cell growth and immune response in DJ-1-dependent glial pathologies in glioblastoma and PD. In models of PD, DJ-1 level dependent phenotypes can be rescued by antioxidant treatment that reduces the GSSG/GSH ratio and ROS levels.

## Introduction

Astrocytes are the most abundant glial subtype in the brain and are critical for the normal functioning of the brain. Recently, there has been increasing evidence for an important role of astrocytes in the pathogenesis of PD^1^. Under physiological conditions, one of the main functions of astrocytes is to metabolically support the surrounding neurons. Astrocytes have many different neuroprotective functions like releasing neurotrophic factors, producing antioxidants like glutathione (GSH), and disposing of neuronal waste products^2^. Since neurons are highly energy demanding cells^3^, it is crucial that astrocytes support their metabolism by providing them with glutamine, which can be converted into glutamate and fueled into the neuronal tricarboxylic acid cycle (TCA)^4^. Astrocytes are also involved in inflammatory response or promote immunosuppression and tissue repair. In pathological conditions like PD, astrocytes produce inflammatory cytokines^5^. In fact, astrogliosis is a common pathological feature in PD^6,7^. In addition, impaired astrocytes contribute to PD-linked pathological mechanisms like oxidative stress, neuroinflammation, and mitochondrial impairment^2^. Therefore, targeting astrocytic dysfunction to repair their neuroprotective ability may represent a therapeutic approach to prevent progressive neurodegeneration in PD.

The *PARK7* gene, encoding the protein DJ-1, was initially described as an oncogene as it was isolated in the course of screening for c-Myc-binding proteins in 1997^8^. In 2003, a large deletion and missense mutation in *PARK7* was identified in Italian and Dutch PD patients, leading to the discovery of *PARK7* as a causative gene for familial PD with recessive inheritance^9^. DJ-1 is localized in the cytoplasm, nucleus and mitochondria^10^. In the human brain, DJ-1 displays much higher expression levels in astrocytes than neurons^11^. In PD patient-brains, it was shown that DJ-1 is increased in reactive astrocytes^12^. Several studies in mice showed that DJ-1 overexpression in astrocytes resulted in protection from parkinsonism due to rotenone-induced neuronal cell death and that DJ-1 knockdown or knockout impaired the neuroprotective capacity of astrocytes and decreased neuronal survival^13^. These observations indicate an important role of DJ-1 for astrocytic function and astrocyte-mediated neuronal protection. There is increasing evidence for a direct involvement of DJ-1 in cellular energy metabolism via effects on glycolysis and the TCA cycle^14^. So far, it is not known how DJ-1 affects astrocytic and neuronal metabolism in PD. Astrocytes are the major route for brain glucose uptake during periods of strong synaptic activity, indicating that astrocytic glucose uptake is of key importance to neurons^15^. The loss of DJ-1 in astrocytes could therefore also influence neuronal metabolism and viability. On the other hand, overexpression of DJ-1 could enhance brain metabolism and enable increased cell proliferation. Epidemiologic studies have shown that there is an inverse correlation for gene expression of Parkinson’s disease (PD) and cancer associated genes. Genes that are downregulated in PD are upregulated in cancer and *vice versa*. In fact, high expression of *PARK7* is associated with high grade and poor prognosis in glioma patients due to its influence on cell cycle and apoptosis^16^. Over the last years, emerging evidence indicated astrocytes as cells of origin for GBM^17,18^.

In this study, we analyzed the effects of DJ-1 deficiency or overexpression in patient-derived induced pluripotent stem cell (iPSC) astrocytes and GBM cell lines. We show that metabolic activity, cell growth and immune response upon IL-1β stimulation in models of PD and GBM were dependent on DJ-1 levels. We postulate that the metabolic involvement of DJ-1 in glial cells of models of PD and GBM represents the basis for a better understanding of neurodegeneration.

## Results

### Astrogliosis in DJ-1 PD patient brain

In 2016, Taipa and colleagues described a case of a homozygous DJ-1-mutant patient (p.L172Q mutation) with early-onset parkinsonism with first symptoms at the age of 22. The mutation in the *PARK7* gene causes DJ-1 protein loss and the patient brain showed diffuse Lewy body and astrogliosis^19^. To further analyze astrogliosis in this patient, we obtained sections from the cortex (of the patient, male, 49 years old, and a gender-matched control, male, 47 years old). We found that the patient brain showed a higher abundance of the activated astrocyte marker GFAP when compared to the control, which was accompanied by enlarged cell bodies and processes, known indicators for astrocyte activation^20^ (Figure 1A). In addition, astrocytes in the patient brain displayed reduced intensity of staining for the astrocyte marker aldolase c (Aldoc), a glycolytic enzyme (Figure 1B), implying that the patient astrocytes shift from a metabolically active to a reactive state (more GFAP staining). In addition, the microglial marker Allograft inflammatory factor 1 (Iba1) showed an increased staining in the cortex of the DJ-1 patient when compared to the control, indicating increased neuroinflammation in the patient (Figure 1C, Suppl. file 1).

**Figure 1:**
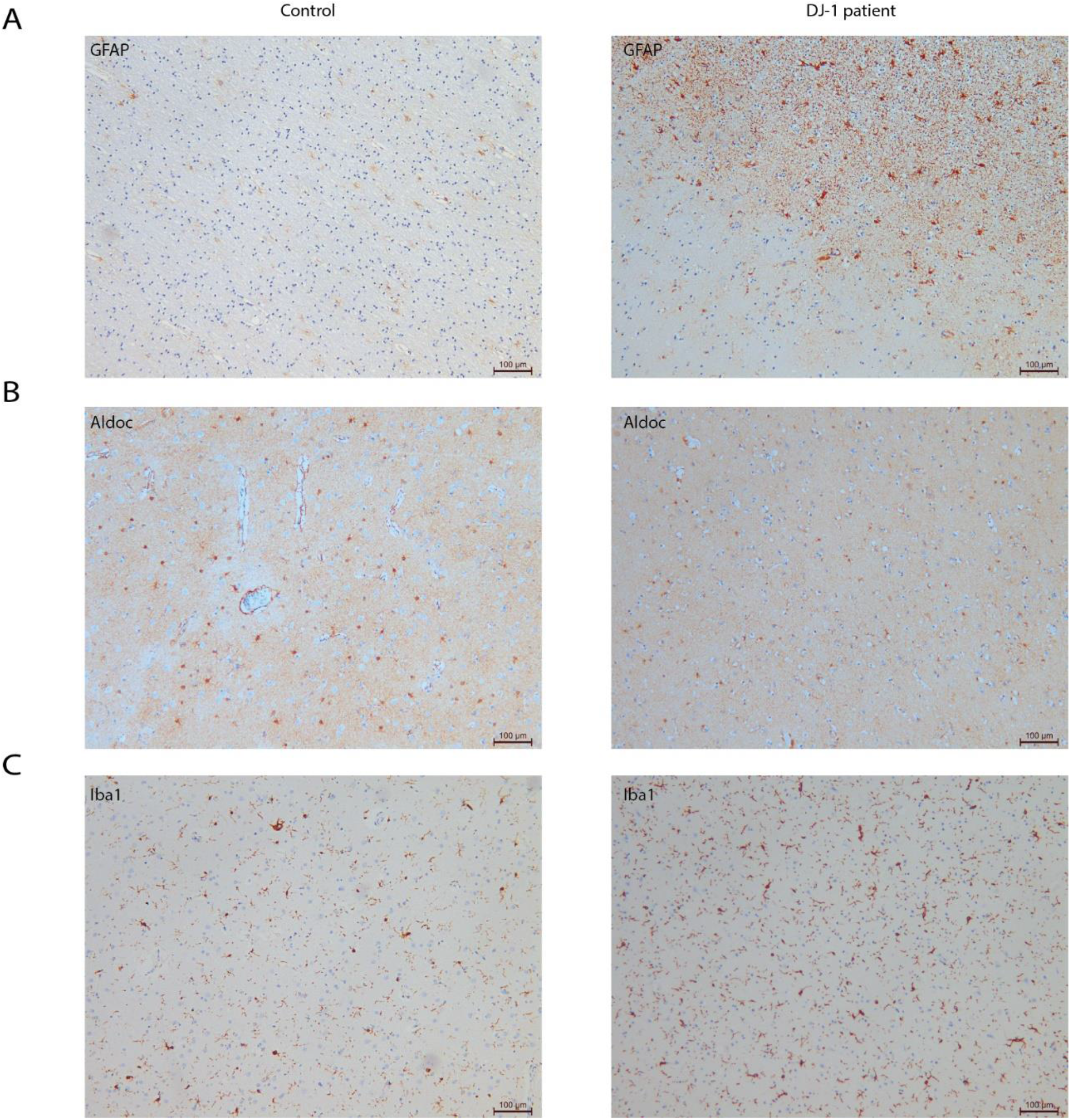
Astrogliosis in DJ-1 patient brain. Cortex section of a healthy donor (control) and the PD patient (DJ-1 patient) stained for **A:** the activated astrocyte marker GFAP. **B:** the astrocyte marker aldolase c (Aldoc). **C:** the microglial marker Allograft inflammatory factor 1 (Iba1).

### Increased immune response in DJ-1 deficient astrocytes

To further investigate the effect of DJ-1 deficiency in PD astrocytes, we derived astrocytes from induced pluripotent stem cells (iPSCs) carrying two different PD-associated DJ-1 mutations. We used isogenic pairs with two different homozygous DJ-1 mutations - P158Δ in-frame deletion (DelP) and DelP gene-corrected (GC), and c.192G>C (C4 mut) and C4, respectively^21–23^. Sixty days-old astrocytes were used for all experiments. The cells showed an astrocytic morphology, and the majority stained positive for canonical astrocyte markers like GFAP, S100b, Vimentin, EAAT2, NFIA, ID3, and EZRIN. No neuronal contamination was observed, as assessed by markers for TUJ1 and MAP2 (Suppl. Fig. 1). Expression of typical markers of astrocyte precursors is shown in supplementary figures (Suppl. Fig. 2-3). Gene expression of all cell lines was assessed using next generation RNA sequencing (Suppl. Fig. 4 and methods). Differential expression analysis showed downregulation of 146 genes and upregulation of 56 genes in C4 mut astrocytes compared to isogenic control C4 astrocytes using a with log2-fold change cut-off of ± 1 with an adjusted p value < 0.05 (Suppl. Fig. 5A and B). Ingenuity pathway analysis (IPA)^24^ on the differentially expressed genes^24^ considering −log(p-value) > 2 identified that the highest ranked upregulated pathway in DJ-1-deficient astrocytes was neuroinflammation (z-score = 0.632) (Suppl. Fig. 5C). The graphical summary of this IPA core analysis illustrates the relation of upregulated Interleukin 1 beta (IL-1β) and interferon gamma (IFNG) signaling, immune signaling involved in T cell cytotoxicity and cancer immunotherapy, and apoptosis (Figure 2A and B). Therefore, we first validated the increased immune response in DJ-1-deficient astrocytes, which showed increased gene expression of *C-C Motif Chemokine Ligand 5 (CCL5*), also known as RANTES, upon stimulation with IL-1β for 2 to 12 hours, and for 24 and 48 hours when compared to isogenic controls (Suppl. Fig. 6A). A similar phenotype was observed in DJ-1-deficient microglia (Suppl. Fig. 6B). The increased gene expression levels of *CCL5* in DJ-1-deficient astrocytes resulted in increased secretion of CCL5 protein into the medium (Figure 2C). Concordantly with the increased cytokine expression and release, DJ-1-deficient astrocytes attracted more CD4+ and CD8+ human T cells, as assessed by T cell migration assay, and CCL5 secretion was also increased during the assay after 48 hours upon stimulation with IL-1β (Figure 2D). The migration towards DJ-1-deficient astrocytes was even increased upon knockdown of DJ-1 in control T cells (60% knockdown, see Suppl. Fig. 7), a scenario that is closer to the situation of a PD patient with homozygous DJ-1 mutations in which the T cells are also DJ-1-deficient. Additionally, DJ-1 deficiency also significantly decreased astrocytic proliferation (Figure 2E).

**Figure 2:**
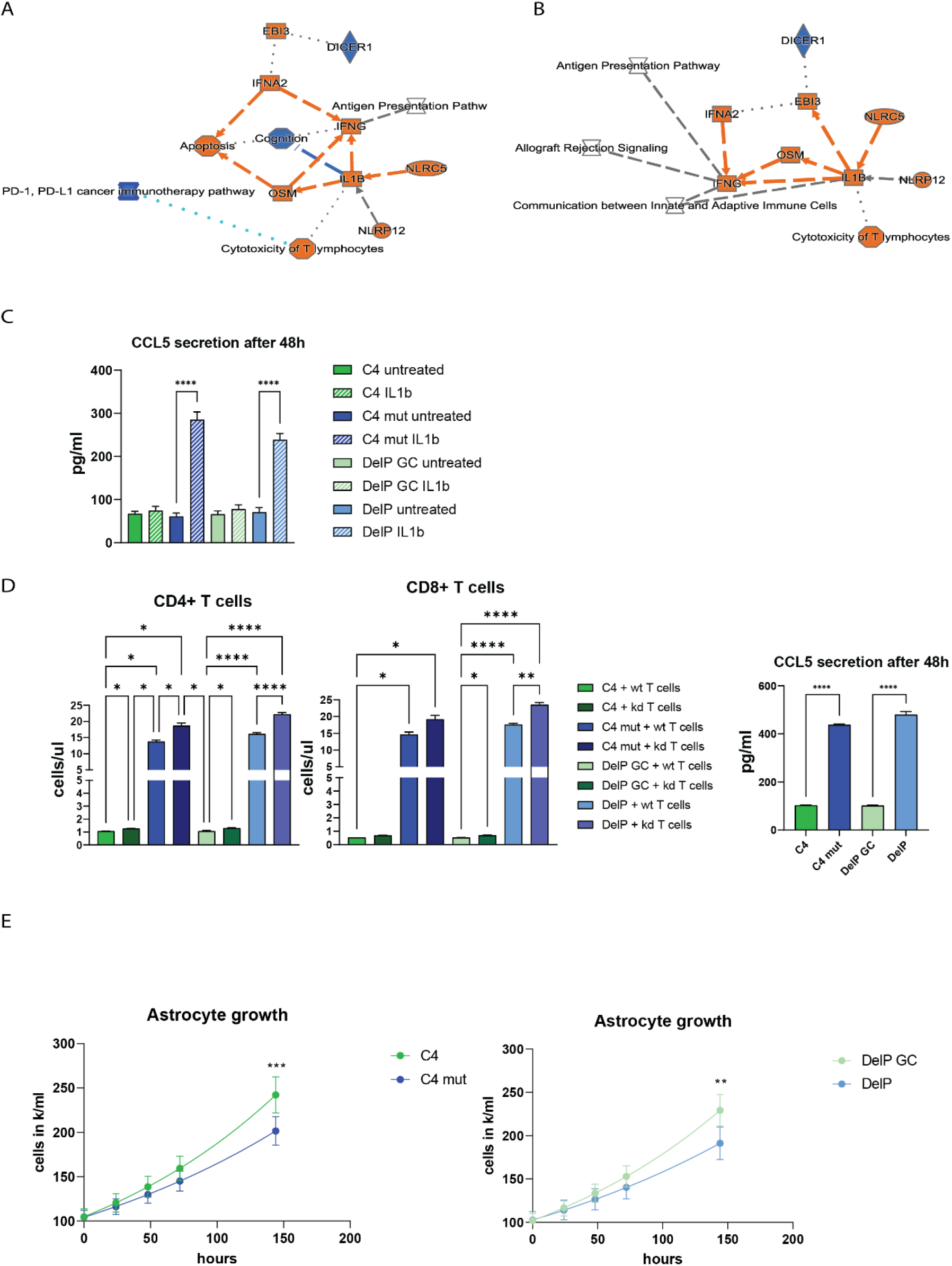
Increased immune response in DJ-1-deficient astrocytes. **A and B:** Ingenuity pathway analysis. N=3. Relaxed cut offs were chosen due to little number of differentially expressed genes cutoff -0.5-0.5 log2foldchange p adj. 0.1. **C:** CCL5 secretion. N=3. Error bars show SEM. One-way ANOVA was used with Tukey’s multiple comparisons. **D:** T cell migration towards astrocytes and CCL5 secretion measured in these assays. N=3. Error bars show SEM. One-way ANOVA was used with Tukey’s multiple comparisons. **E:** Astrocyte growth (N=3) with SEM error bars. Non-linear fit was calculated (Exponential Malthusian growth) and 2-way ANOVA for each time point with Tukey’s multiple comparisons. Significance is indicated for the last time point. **D-F:** p <0.0001 = ****, p <0.001 = ***, p <0.01 = **, p <0.05 = *.

### DJ-1 deficiency in astrocytes causes impaired metabolic carbon contribution and GSH levels

Based on the observed decrease in cell growth, we next investigated whether DJ-1-deficient astrocytes might have an impairment in energy metabolism. We analyzed glucose and glutamine metabolism in more detail by performing glucose and glutamine tracing. DJ-1-deficient astrocytes took up less glucose and released less lactic acid (Figure 3A)^25^. In line with reduced glucose uptake, glucose carbon contribution to the TCA was decreased (Figure 3B). However, the glycolytic carbon contribution towards 3PG and lactate was unaffected, as seen by analyzing the production of pyruvate from 3PG (by calculating the ratio of M3 pyruvate over M3 3PG) and the ratio of M3 lactate over M3 pyruvate (Figure 3C). Pyruvate entry into the TCA was reduced as assessed via the analysis of the production of M2 citrate from M3 pyruvate (Figure 3D). Consequently, the TCA cycling was reduced due to decreased production of M4 citrate from M2 citrate (Figure 3E). Labelling of glutamate was decreased indicating a deficiency of the cell to provide glutamate and eventually glutamine via glucose metabolism (Figure 3B). Glutamate is used by the cell for the synthesis of glutathione (GSH), the most important molecule in cellular oxidative stress response, and like lactic acid, glutamine is released by astrocytes to metabolically support neurons^4^. The reduced metabolic carbon contribution seen in deficient astrocytes by metabolic tracing was functionally confirmed by measurements of the extracellular acidification rate (ECAR) (Figure 3F) and the oxygen consumption rate (OCR) (Figure 3G), which revealed decreased glycolysis and oxidative phosphorylation (OXPHOS) compared to controls (Figure 3F and G). The decreased contribution of glucose to the TCA and glutamate production raised the question about the ability of the cell to compensate via the use of glutamine. Glutamine tracing revealed that glutamine uptake was significantly increased in DJ-1 -deficient astrocytes, and that over 95% of the glutamine detected in all cell lines was taken up from the medium (over 95% carbon contribution, Figure 3H). Yet, glutamine carbon contribution to all TCA cycle metabolites was decreased in DJ-1-deficient astrocytes compared to isogenic controls (Figure 3H). Interestingly, DJ-1 deficient astrocytes used more glutamine for fueling into the TCA, as seen by the increased ratio of M5 alpha-ketoglutarate over M5 glutamate (Figure 3I). However, increased alpha-ketoglutarate labeling did not translate into increased succinate labeling, as the production of succinate from alpha-ketoglutarate was significantly decreased (Figure 3J). Glutamine carbon contribution of the subsequent TCA cycle metabolites and the cycling of the TCA were therefore also significantly reduced in DJ-1 deficient astrocytes (Figure 3K). This means that the increased glutamine carbon contribution to alpha-ketoglutarate is not channeled towards the TCA cycle to compensate for lower glycolytic contribution. Glutamine can also be used for GSH synthesis, which is known to be decreased in DJ-1-deficient cells^26^. Indeed, GSH and oxidized glutathione (GSSG) labeling was increased compared to isogenic controls (Figure 3L and 3M). However, despite increased use of glutamine for GSH production, total GSH levels were decreased and GSSG levels still increased in DJ-1-deficient astrocytes when compared to isogenic controls, as assessed by luminescence-based quantification (Figure 3N). As more glutamine from the medium was used by DJ-1-deficient astrocytes to synthesize GSH, we next analyzed whether doubling the amount of glutamine in the medium could rescue GSH levels. Upon high glutamine supplementation (4 mM), GSH levels in the DJ-1-deficient astrocytes reached wildtype levels (Figure 3N). GSSG levels remained increased in DJ-1-deficient astrocytes when compared to the isogenic counterparts, indicating that the increased GSH levels upon glutamine supplementation are being used to diminish increased ROS levels (Figure 3N). GSSG/GSH ratio was significantly decreased upon glutamine supplementation (Figure 3N), which indicates a decrease in oxidative stress upon glutamine supplementation in DJ-1-deficient cells, but not in the isogenic controls.

**Figure 3:**
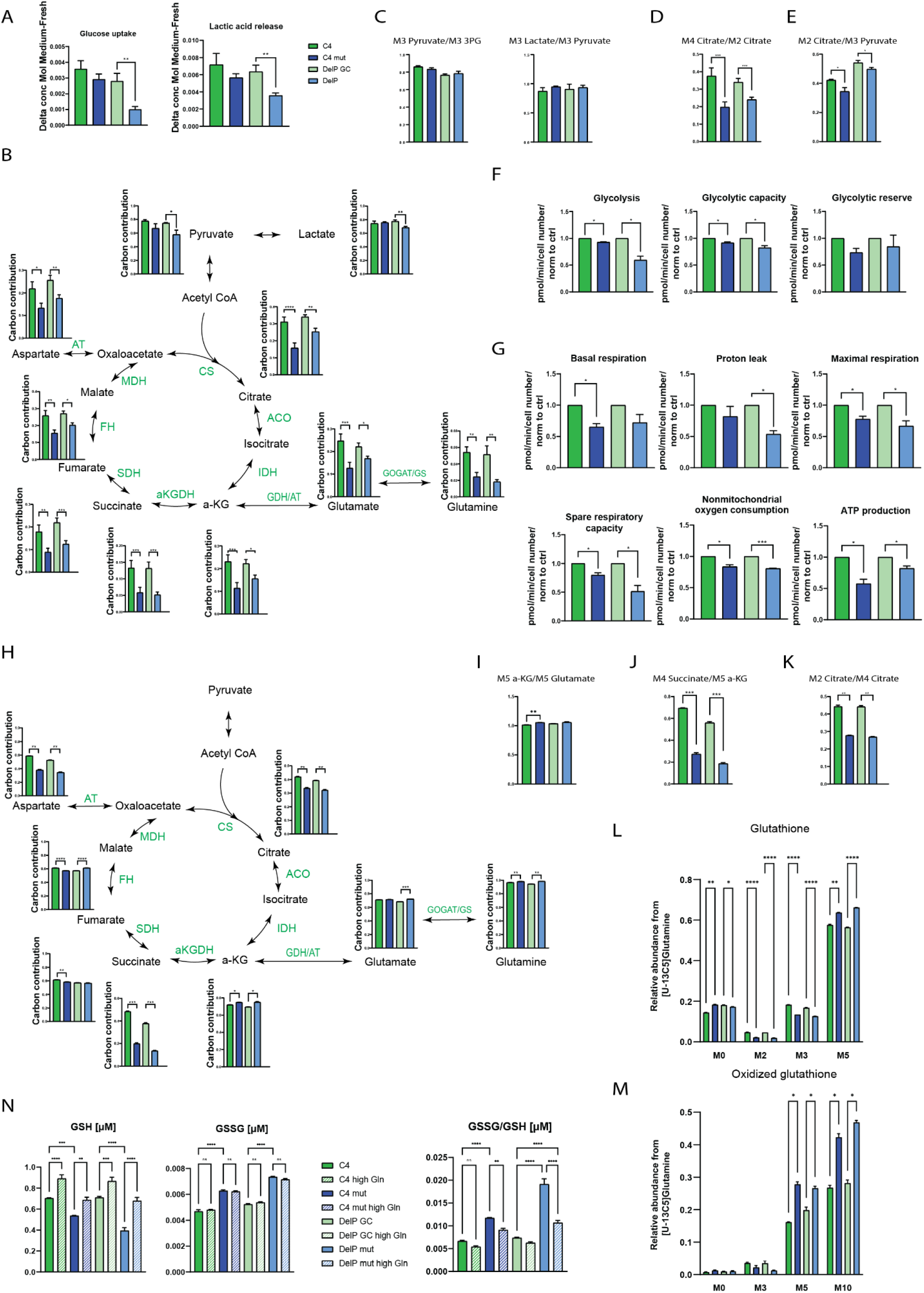
DJ-1 deficiency in astrocytes causes impaired metabolic carbon contribution and GSH levels. **A:** Analysis of uptakes and release rates from the medium by GC-MS. N=3-5. Error bars show SEM. Two-tailed paired T test was used. **B:** Analysis of glucose metabolism using [U-13C6]Glucose tracing. N=3-5. ^13^C incorporation in metabolites was analyzed by GC-MS, resulting in heavier metabolites (M1+x), whereas no ^13^C incorporation corresponds to M0. The graphs show the carbon contribution for each metabolite (calculation see methods part). Error bars show SEM. Two-tailed paired T test was used. **C-E:** Ratios: The higher the ratio, the more production of the respective metabolite from its precursor. Error bars show SEM. Two-tailed paired T test was used. N=3-5. **F-G:** Extracellular flux analysis using Seahorse. N=3-5. Error bars show SEM. Two-tailed paired T test was used. **H-M:** [U-13C5]Glutamine tracing. N=3. Error bars show SEM. Paired T test was used, for GSH and GSSG 2-way ANOVA with Turkey’s multiple comparisons. **N:** GSH and GSSG measurement in cell lysates. N=3. Error bars show SEM. ANOVA was used. **N:** GSH and GSSG level assessed by luminescence-based quantification. N=3. Error bars show SEM. One-way ANOVA was used with Tukey’s multiple comparisons. **A-N:** p <0.0001 = ****, p <0.001 = ***, p <0.01 = **, p <0.05 = *.

### Increased glutamine supply reduces ROS levels and rescues immune phenotype and growth impairment of DJ-1 deficient astrocytes

DJ-1 is a known ROS scavenger^26–29^ and loss of DJ-1 protein was shown to increase ROS in mouse primary astrocyte cultures^30–34^. Consistent with these findings, DJ-1-deficient human astrocytes had elevated levels of cellular ROS, RNS and superoxide, which are predominantly produced as by-products of mitochondrial respiration (Figure 4A). The observed increase in GSSG levels in DJ-1 deficient cells (Figure 3N) indicates failure in oxidative stress response. We saw that glutamine uptake was increased in DJ-1-deficient astrocytes to produce GSH (Figure 3L). Thus, we further assessed whether increasing the amount of glutamine could also rescue the increased ROS levels. ROS levels were significantly reduced by high glutamine treatment in all cell lines, but not as efficiently as when treated with the positive control N-Acetyl-Cysteine (NAC) (Figure 4B and C). ROS are important signaling molecules involved in immune signaling^35^. Therefore, we analyzed whether high glutamine or NAC supplementation in the astrocyte cultures could rescue immune related phenotypes. Both treatments reduced cytokine release (Figure 4D) and T cell migration (Figure 4E). Furthermore, doubling the amount of glutamine in the astrocyte medium rescued astrocyte cell growth of DJ-1-deficient astrocytes compared to isogenic controls (Figure 4F). The dependency on glutamine uptake can also be seen in cell survival rates. In baseline conditions, early apoptosis was significantly increased in DJ-1 deficient astrocytes compared to isogenic controls. Additional glutamine starvation for 4 hours further led to a significant increase of apoptosis in DJ-1-deficient astrocytes, which was not observed in isogenic controls (Figure 4G).

**Figure 4:**
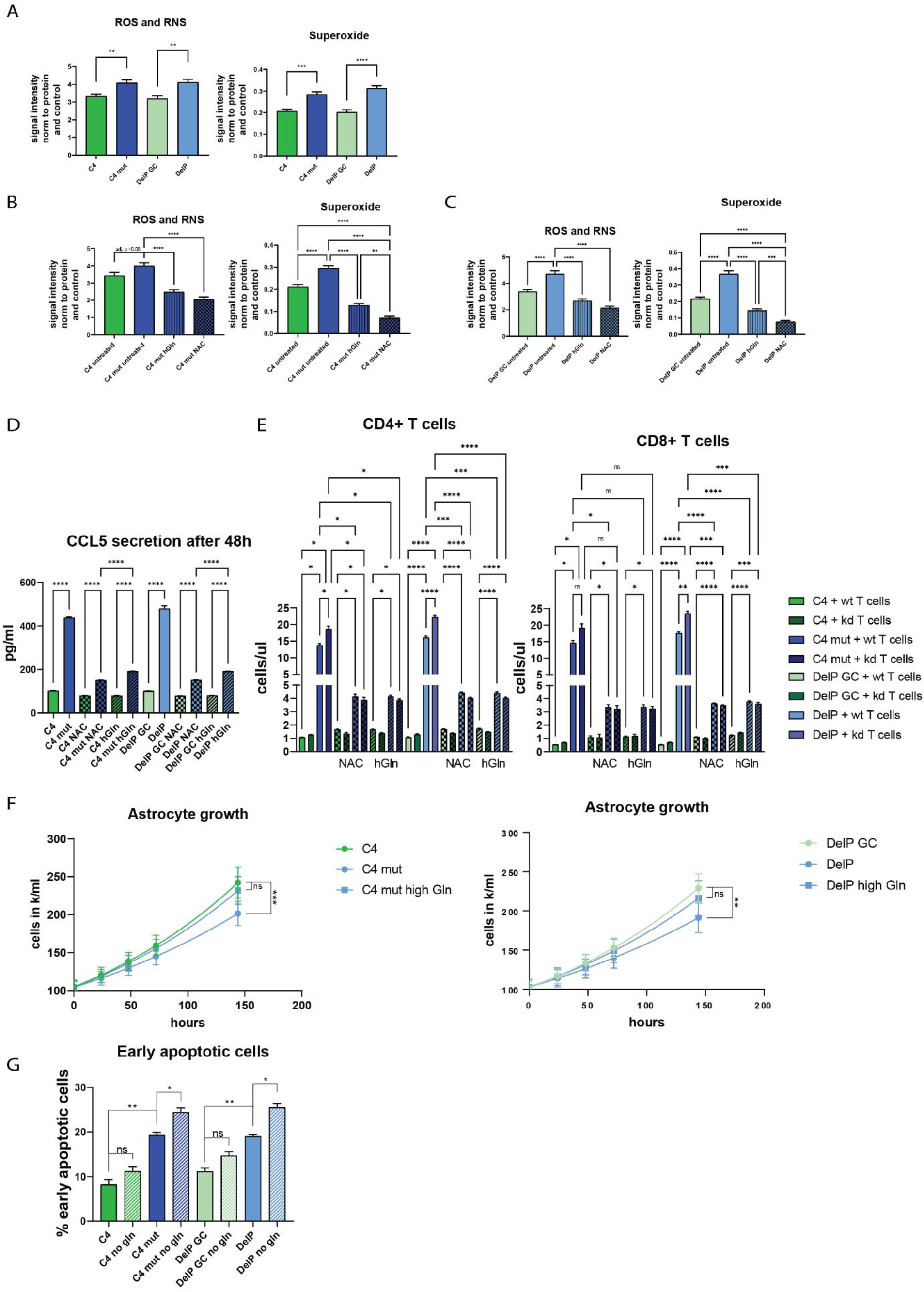
Increased glutamine supply reduces ROS levels and rescues cellular phenotypes. **A:** Cellular ROS/RNS (left) and superoxide (right) levels in astrocytes. N=3. Error bars show SEM. Two-tailed paired T test was used. **B-C:** Cellular ROS/RNS (left) and superoxide (right) levels in astrocytes upon high glutamine or NAC supplementation. N=3. Error bars show SEM. One-way ANOVA with Šídák’s multiple comparisons test was used. **D:** CCL5 secretion upon high glutamine or NAC supplementation. Error bars show SEM. One-way ANOVA was used with Tukey’s multiple comparisons. **E:** T cell migration towards astrocytes. N=3 (3 different blood donors with 3 independent astrocyte differentiations). Error bars show SEM. One-way ANOVA was used with Tukey’s multiple comparisons. **F:** Astrocyte growth (N=3) with SEM error bars. Non-linear fit was calculated (Exponential Malthusian growth) and 2-way ANOVA for each time point with Tukey’s multiple comparisons. Significance is indicated for the last time point. **G:** Early apoptosis assessed by Annexin V staining. N=3. Error bars show SEM. paired T test was used. **A-G:** p <0.0001 = ****, p <0.001 = ***, p <0.01 = **, p <0.05 = *.

### Metabolic and growth impairment are reversed in DJ-1 overexpressing astrocytes and GBM cell lines

Our observation that DJ-1 deficiency negatively modulates astrocyte metabolism and growth while increasing immune response led us to hypothesize that these mechanisms might be inversely regulated in GBM cases due to increased DJ-1 levels in GBM. Therefore, we generated DJ-1-overexpressing iPSC-derived astrocytes as an oncogenic model to study the influence of elevated DJ-1 levels in astrocytes. Lentiviral overexpression was established in the iPSC line C4 to keep the same genetic background as the isogenic pair used to model PD (Suppl. Fig. 8 and 9). To study the role of DJ-1 in GBM, we used three different GBM cell lines, namely U251, U87 and LN229. All three lines displayed upregulated DJ-1 protein levels when compared to C4 astrocytes (Suppl. Fig. 10A).

Gene expression of DJ-1-overexpressing and C4 astrocytes was profiled using RNA sequencing (Suppl. Fig. 4). Differential expression analysis revealed that 323 genes were downregulated and 225 upregulated in DJ-1-overexpressing astrocytes compared to isogenic control C4 with log2-fold change of -1 and padj of 0.05 (Suppl. Fig. 11). IPA gene expression analysis showed an upregulation of pathways involved in cell cycle control, cell growth, and especially pathways associated with proliferation of tumor cells (Figure 5A and B) compared to C4 astrocytes. Thus, we again analyzed the growth behavior of the cells. Concordantly, C4 DJ-1 overexpression astrocytes showed increased cell growth compared to C4 control astrocytes (Figure 5B). On the other hand, cell growth was reduced upon DJ-1 knockdown in the GBM cells (Figure 5C), which could be rescued by doubling the amount of glutamine in the medium (Figure 5D).

**Figure 5:**
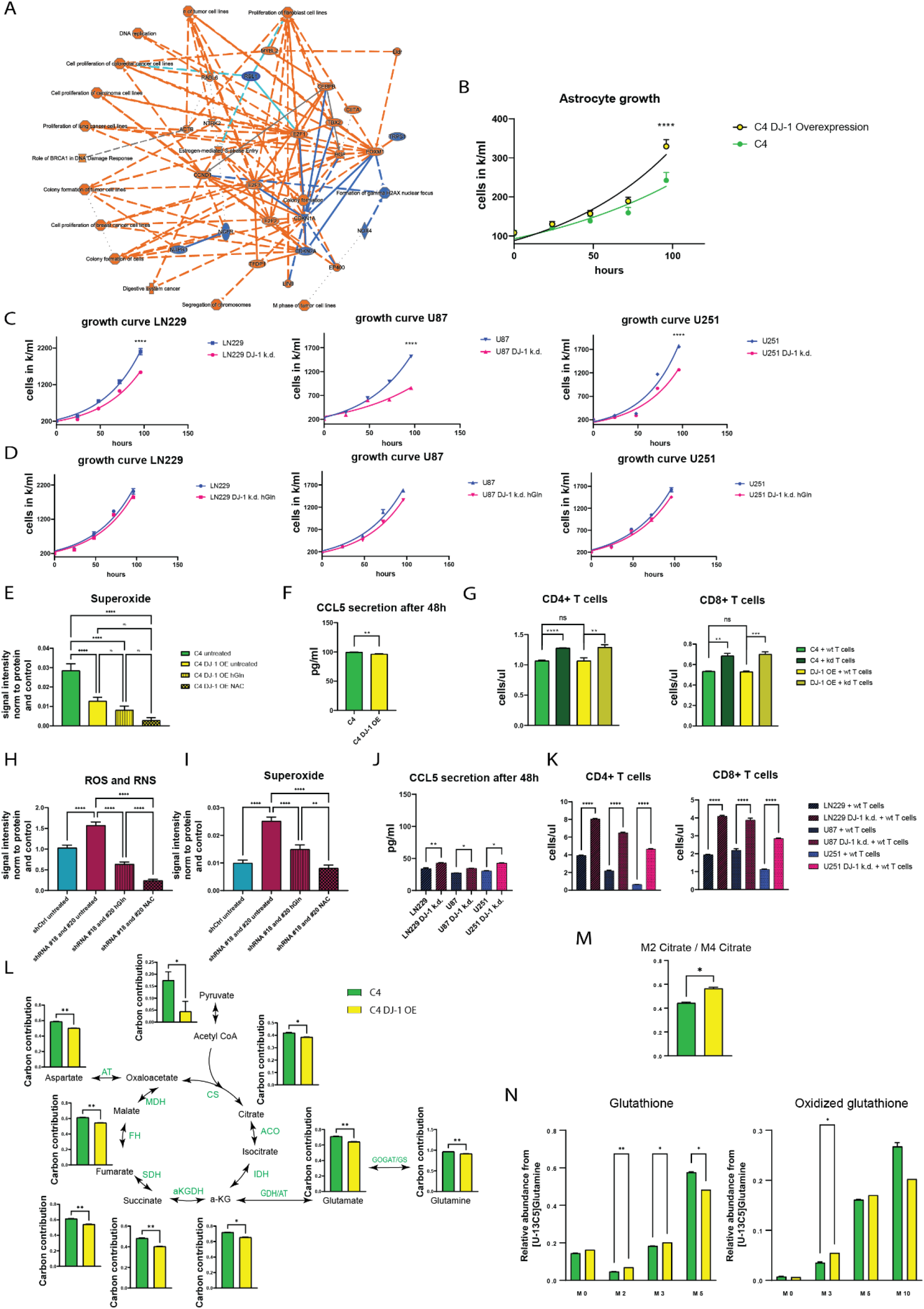
Growth impairment is reversed in DJ-1 overexpressing astrocytes and GBM cell lines. **A:** Ingenuity pathway analysis of DJ-1 overexpression astrocytes versus wildtype astrocytes. N=3. Upregulated pathways are shown in orange, downregulated ones in blue. **B:** Astrocyte cell growth with SEM error bars. N=3.Nonlinear fit was calculated with exponential (Malthusian) growth. 2-way ANOVA for each time point with Tukey’s multiple comparisons. Significance is indicated for the last time point. **C-D:** Growth curve of GBM cell lines. N=3. Nonlinear fit was calculated with exponential (Malthusian) growth. 2-way ANOVA for each time point with Tukey’s multiple comparisons. Significance is indicated for the last time point. **E**: Mitochondrial ROS level. N=3. Error bars show SEM. One-way ANOVA was used with Tukey’s multiple comparisons. **F:** CCL5 secretion of astrocytes assessed by ELISA. Error bars show SEM. Two-tailed paired T test was used. **G:** T cell migration towards astrocytes. Error bars show SEM. One-way ANOVA was used with Tukey’s multiple comparisons. **H-I:** ROS levels in GBM cells. The mean of the 3 lines is shown. Error bars show SEM. One-way ANOVA was used with Tukey’s multiple comparisons. **J:** CCL5 secretion of GBM cells assessed by ELISA. Each line is shown separately. N=1. Error bars show SEM. One-way ANOVA was used with Tukey’s multiple comparisons. **K:** T cell migration assay in GBM cell lines. Each line is shown separately. N=1. Error bars show SEM. One-way ANOVA was used with Tukey’s multiple comparisons. **L:** Glutamine LC-MS tracing in astrocytes with DJ-1 overexpression compared to wildtype astrocytes. N=3. Error bars show SEM. Two-tailed paired T test was used. **M:** Glutamine LC-MS tracing-based measurement. Ratio M2 citrate over M4 citrate. **N:** Glutamine LC-MS tracing-based measurement of GSH and GSSG. N=3. Error bars show SEM. Two-tailed paired T test was used. **B-M:** p <0.0001 = ****, p <0.001 = ***, p <0.01 = **, p <0.05 = *.

DJ-1 overexpression in astrocytes caused reduced ROS levels, which were further reduced by high glutamine and NAC supplementation (Figure 5E). The overexpression decreased CCL5 secretion (Figure 5F), but did not affect T cell migration (Figure 5G). On the other hand, the knockdown of DJ-1 in the GBM cells increased ROS levels (Figure 5H and I) and led to increased CCL5 secretion (Figure 5J) and consequently T cell migration (Figure 5K). Taken all together, overexpression of DJ-1 in astrocytes lead to opposite cellular phenotypes than observed in DJ-1-deficient astrocytes. However, glutamine carbon contribution to the TCA (Figure 5L and M) and to GSH production (Figure 5N) was decreased in DJ-1 overexpression astrocytes compared to wildtype astrocytes. This suggests that low ROS levels and a sufficient metabolic activity make the cells less dependent on glutamine. In that case an increased carbon contribution of glucose to the TCA cycle should be seen.

Indeed, when analyzing glucose metabolism using labeled glucose, DJ-1 overexpression in astrocytes increased the carbon contribution from glucose to the TCA cycle (Figure 6A). Expectedly, we saw that knockdown of DJ-1 in the three different GBM cell lines reduced the carbon contribution from glucose to the TCA cycle and the reduction correlated with the knockdown efficiency of the three different shRNAs on protein levels in the GBM lines (Figure 6A and B). This suggests a dependency of the metabolic activity on DJ-1 levels in both cell types. Importantly, glutamate is known to be crucial for cancer cell metabolism^36,37^. The contribution of glucose to its labeling was significantly increased in the oncogenic model, and DJ-1 knockdown in the GBM cells significantly reduced the carbon contribution of glucose to glutamate. Analysis of OXPHOS activity by OCR measurements revealed that the increased contribution from glycolysis in DJ-1 overexpressing astrocytes did not lead to an increase in mitochondrial respiration (Figure 6C). However, DJ-1 knockdown in GBM cells significantly decreased OXPHOS suggesting that increased DJ-1 levels are not sufficient to increase mitochondrial respiration but are essential to maintain an increased metabolic activity.

**Figure 6:**
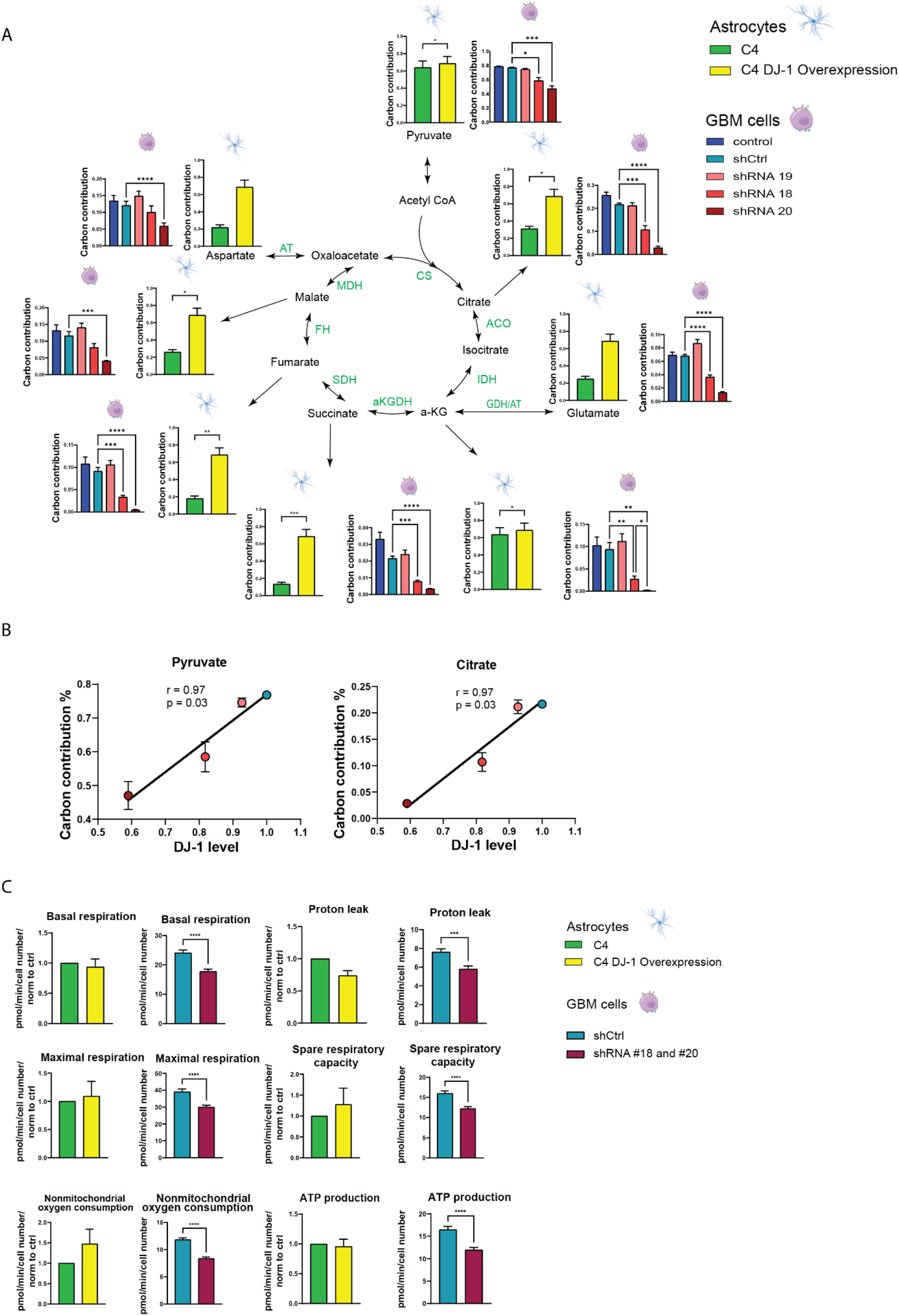
Metabolic impairment is reversed in DJ-1 overexpressing astrocytes and GBM cell lines. **A:** Analysis of glucose metabolism using [U-13C6]Glucose tracing. Error bars show SEM. One-way ANOVA was used with Tukey’s multiple comparisons. **B:** Analysis of correlation of DJ-1 protein levels and carbon contribution using simple linear regression. **C:** Extracellular flux analysis of oxygen consumption rate in astrocytes and GBM cells. Mean of the 3 different GBM cell lines is shown. Error bars show SEM. Paired T test was used. **A-C:** p <0.0001 = ****, p <0.001 = ***, p <0.01 = **, p <0.05 = *.

### NAC supplementation rescues TCA cycle carbon contribution deficits in DJ-1 deficient astrocytes and high glutamine supplementation increases TCA cycling and GSH synthesis from glutamine

To understand whether the observed phenotypes of increased immune response and impaired cellular metabolism were caused by decreased GSH levels and subsequently increased ROS in the absence of DJ-1, we then analyzed glucose and glutamine metabolism in the patient-derived isogenic astrocytes by LC-MS tracing after treating the cells with NAC as a positive control or doubling the amount of glutamine in the medium. NAC supplementation rescued the glucose carbon contribution to all TCA intermediates (Figure 7A). The increased TCA carbon contribution was not only due to increased carbon contribution of glucose to pyruvate but also due to increased pyruvate entry into the TCA cycle (Figure 7B). Consequently, glucose carbon contribution to glutamate was brought back to wildtype levels (Figure 7A). In contrast, high glutamine did not rescue glucose carbon contribution to TCA cycle intermediates (Figure 7A). Glutamine tracing revealed that 99% of the intracellularly detected glutamine in both cell lines and all conditions was taken up from the medium and converted to glutamate (Figure 7C). Increased fueling of the TCA with glutamate to produce alpha-ketoglutarate was detected in case of DJ-1 deficiency (Figure 7C, D). However, only NAC treatment rescued the TCA cycle intermediate labeling (Figure 7C) to wildtype levels, as the conversion of alpha-ketoglutarate to succinate was rescued in DJ-1 deficient astrocytes (Figure 7E). In contrast to NAC treatment, supplying the cells with 4 mM glutamine did not rescue the glutamine carbon contribution to the TCA cycle intermediates (Figure 7C). The analysis of the ratio of M4 succinate over M5 alpha-ketoglutarate showed that the additionally provided glutamine did not enable the cells to use it efficiently for the TCA and to compensate for the reduced glucose carbon contribution to the TCA (Figure 7E). However, and confirming the results from earlier experiments (Figure 3L-N), the increase of glutamine concentration led to an increased carbon contribution from glutamine to GSH (Figure 7F, M5 GSH) compared to wildtype cells and to the same cells in basal conditions or treated with NAC. NAC supplementation significantly decreased carbon contribution from glutamine to GSSG, but high glutamine did not affect GSSG labeling (Figure 7G). This suggests that additionally supplied glutamine is used for oxidative stress response, but that the protective effect is not as strong as treatment with NAC that caused a stronger reduction of ROS levels, especially of superoxide (Figure 4B and C) and hence, was able to not only rescue the growth and immune phenotypes, but also the TCA cycle insufficiency.

**Figure 7:**
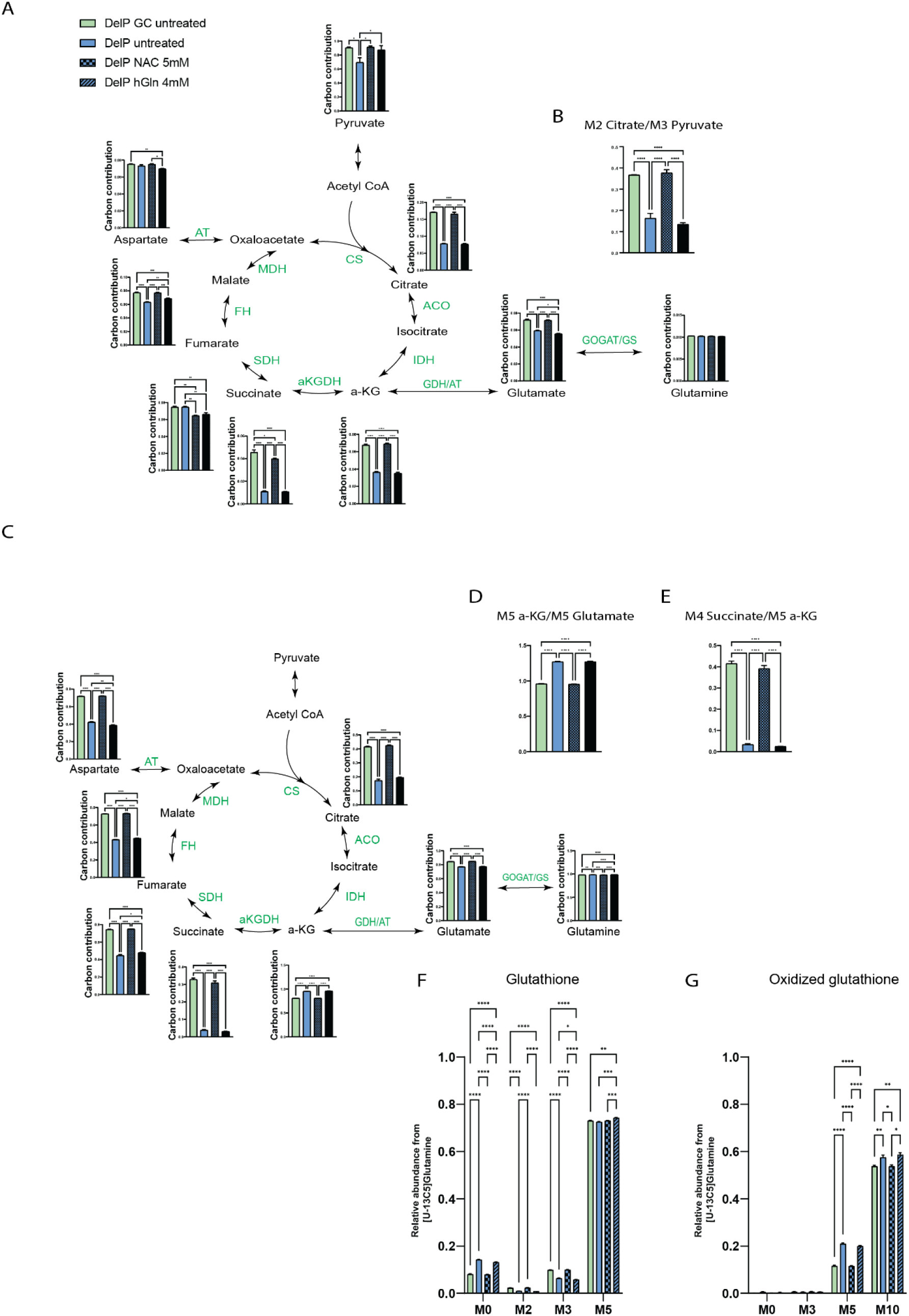
NAC supplementation rescues TCA cycle carbon contribution deficits in DJ-1 deficient astrocytes. **A:** Analysis of glucose metabolism using [U-13C6]Glucose tracing. N=3. 13C incorporation in metabolites was analyzed by GC-MS, resulting in heavier metabolites (M1+x), whereas no ^13^C incorporation corresponds to M0. The graphs show the carbon contribution for each metabolite (calculation see methods part). Error bars show SEM. One-way ANOVA was used with Tukey’s multiple comparisons. **B:** Ratios for glucose tracing. N=3. Error bars show SEM. One-way ANOVA was used with Tukey’s multiple comparisons. **C-G:** [U-13C5]Glutamine tracing results. N=3. Error bars show SEM. One-way ANOVA was used with Tukey’s multiple comparisons. **A-G:** p <0.0001 = ****, p <0.001 = ***, p <0.01 = **, p <0.05 = *.

## Discussion

Astrocytes play an important role in the pathogenesis of PD, and GBM is claimed to be originating from astrocytes^17,381^. Thus, we analyzed the role of DJ-1 downregulation in PD, and DJ-1 upregulation in GBM to shed light on the opposite disease-related phenotypes of PD and GBM associated with diverging DJ-1 levels.

In human cortex of a PD patient with a DJ-1 mutation, we found increased astrogliosis^19^, which supports an activated immune mechanism involving astrocytes in the neurodegenerative process. Increased astrogliosis in PD midbrain was observed in previous studies of human midbrain of patients with idiopathic PD^7^, and neuroinflammation is increasingly recognized to play a critical role in the pathology of PD^392^. Our observation in human astrocytes deficient of DJ-1 was supported by next generation RNA sequencing showing an upregulation of proinflammatory pathways, which was subsequently validated by increased cytokine expression and release upon stimulation. It was previously shown that IL-1β and IL-6 are elevated in the brains from idiopathic PD patients^40^. A recent publication showed an increased cytokine release in human astrocytes derived from a PD patient carrying a G2019S mutation in LRRK2^41^. Astrocytes and microglia from DJ-1-deficient mice showed increased inducible NO synthase (iNOS) levels^42^, which indicates that dysfunctional astrocytes may act as source for neuroinflammation. Kahle and colleagues also demonstrated higher astrocyte reactivity by showing increased cytokine release in astrocytes from DJ-1^-/-^ mice upon lipopolysaccharide (LPS) stimulation compared to controls^30^. Furthermore, astrocytes from DJ-1^-/-^ mice produced more nitric oxide (NO), which was mediated by ROS signaling leading to activation of iNOS^30^. Additional evidence for a primary pathological role of DJ-1^-/-^ astrocytes in neurodegeneration came from *in vitro* studies. Neurotoxicity upon LPS treatment was only observed, when either wildtype or DJ-1^-/-^ neurons were co-cultured with DJ-1^-/-^ astrocytes, but not with wildtype astrocytes^30^. Interestingly, also in the *in vivo* situation of DJ-1 knockout mice no other neurodegeneration was observed as these animals displayed no nigral or striatal loss of dopaminergic neurons and only mild non-motor symptoms^43^, which may indicate that astrocytic pathology did not yet translate into neurodegeneration. Taken together, this strengthens the hypothesis that dysfunctional astrocytes significantly contribute to the pathogenesis of neurodegeneration and precedes neuronal loss *in vivo*. Consistent with an astrocytic activation, we found significantly decreased GSH levels and increased ROS in DJ-1-deficient human astrocytes compared to isogenic controls. ROS are known to regulate the immune response^35,44,45^. In fact, it was shown that mitochondrial ROS can induce proinflammatory cytokine production^46^. A recent study suggests that transiently increased CCL5 expression in mice brains after mild traumatic brain injury (TBI) is a coping mechanism to reduce elevated ROS levels caused by the injury via glutathione peroxidase-1 (GPX1) activation^47^. However, in contrast to the situation after TBI, ROS levels in DJ-1-deficient cells are chronically elevated. In the case of astrocytes, this leads to an increased CCL5 release, which over time can contribute to neuroinflammation^48^. These results are again indicating an important role of astrocytes as direct key players in causing neurodegeneration by maintaining a chronic inflammatory reaction. It was shown that infiltration of CD4+ lymphocytes into the brain contributes to neurodegeneration in a PD mouse model^49^. Here, we assessed the functional relevance of the increased ROS levels and cytokine release in DJ-1-deficient astrocytes and found increased T cell migration towards DJ-1-deficient astrocytes. Previously, it was shown that T cells from DJ-1 KO mice had elevated ROS levels^50^, pointing to the importance of evaluating the effect of lack of DJ-1 in T cells from DJ-1 patients. It is also known that ROS can activate T cells^51^. We observed an even increased T cell migration towards DJ-1-deficient astrocytes after knockdown of DJ-1 in T cells, further implying a contribution to neuroinflammation observed in PD patients with loss of function mutations in the DJ-1 gene.

On the other hand, CCL5 secretion upon stimulation by all three GBM cell lines was lower than in wildtype astrocytes and increased significantly upon DJ-1 knockdown. The knockdown also led to a significant increase in T cell migration towards GBM cells suggesting that elevated DJ-1 levels in GBM might contribute to immune evasion of the tumor. GBM is known for its high metabolic activity^52^ and its immunosuppressive microenvironment that causes T cell dysfunction and therefore impaired T cell migration^53^. These immune evasion mechanisms are decreasing the effectiveness of immune therapy options. Increasing the infiltration of the tumor microenvironment with T lymphocytes is crucial for an improved tumor therapy. The observed increased glycolysis and TCA cycle activity are also connected to T cell exhaustion, as GBM cells deprive the tumor microenvironment from glucose that could be used by infiltrating T cells, which depend on glycolysis^54^. Therapeutic targeting of DJ-1-mediated regulation of cytokine secretion and metabolic modulation could help to decrease tumor growth and immune suppression, which in turn enables effective T cell infiltration, and eventually a better anti-tumor response. On the other hand, for PD, restoring metabolic activity and decreasing cytokine release could counteract neuroinflammation.

Since the T cell migration was reduced upon NAC and glutamine supplementation and cell growth was increased upon glutamine supplementation, we investigated the energy metabolism in DJ-1-deficient astrocytes, which was reduced, as seen by decreased glycolysis, OXPHOS and TCA cycle carbon contribution. We observed the strongest effect of DJ-1 deficiency in astrocytes or DJ-1 knockdown in GBM cells on the TCA flux in the glucose carbon contribution to citrate and glutamine carbon contribution to succinate. These two steps in the TCA are catalyzed by the two rate limiting enzyme complexes pyruvate dehydrogenase complex (PDHC) and alpha-ketoglutarate dehydrogenase complex (OGDHC). While the conversion rate of glucose was not affected during glycolysis up to the stage of pyruvate, our measurements showed a significant decrease of the conversion of pyruvate to citrate. On the other hand, despite an increased glutamine fueling into the TCA until alpha-ketoglutarate in DJ-1 deficient astrocytes, we saw a significantly decreased glutamine carbon contribution to subsequent TCA cycle intermediates. This effect was associated with an impairment in the conversion of alpha-ketoglutarate to succinate. Thus, we observed an inability of astrocytes to use glutaminolysis for efficient energy supply in the absence of DJ-1. However, GSH synthesis from glutamine was increased leading to the conclusion that DJ-1-deficient astrocytes use glutamine for GSH synthesis. Nonetheless, despite increased GSH synthesis, DJ-1-deficient astrocytes had decreased total GSH and increased GSSG levels, which indicates that the increased production from glutamine did not translate into higher total GSH and GSSG levels remained elevated.

Strengthening the rationale of our approach to study underlying pathological mechanisms in rare genetic forms of PD as prototypes for the pathophysiology in idiopathic PD, a recent study also analyzed the metabolism in neuronal precursor cells and dopaminergic neurons derived from induced pluripotent stem cells (iPSCs) of sporadic PD patients^60^. In this study, OGDHC presented a metabolic impediment as its activity was decreased, also resulting in a decreased conversion of alpha-ketoglutarate to succinate causing a reduced TCA cycle flux and thus reduced abundance of citric acid cycle intermediates in these sporadic PD models^60^.

In this study, the hypometabolism in DJ-1 deficient astrocytes resulted in increased apoptosis and decreased cell growth, which was reversed by high glutamine treatment, shifting the increased GSSG/GSH ratio towards increased GSH levels.

It was shown that mitochondrial dysfunction in combination with increased ROS levels cause metabolic failure and hypometabolism, as assessed by metabolomics ([U-13C] glucose tracing) in iPSC-derived astrocytes^62^ of carriers of mutations in the CHMP2B gene involved in frontotemporal dementia (FTD3). Moreover, the study found that the hypometabolism resulted in a switch to a reactive astrocyte phenotype with subsequent increased release of neurotoxic cytokines^62^. Here, we assessed the functional relevance of the ROS-hypometabolism axis in DJ-1-deficient astrocytes and found increased CCL5 secretion and T cell migration towards DJ-1-deficient astrocytes upon stimulation with IL-1β. This effect was reversed by supplying glutathione precursors, indicating that the reduced metabolic activity in the DJ-1 deficient astrocytes is leading to increased astrocytic reactivity attracting more T cells when compared to isogenic counterparts.

Taken together, we found a vicious cycle of reduced GSH levels causing increased ROS levels causing decreased TCA carbon contribution, which in turn leads to reduced production of the GSH precursor glutamine.

In contrast to the observations in DJ-1-deficient astrocytes, DJ-1 overexpression had the inverse effect as it resulted in increased cell growth, metabolic activity and reduced ROS compared to wildtype astrocytes. Ingenuity pathway analysis revealed an upregulation of cancer-associated pathways involved in cell proliferation and cell cycle regulation, which again confirms DJ-1 overexpression in astrocytes as oncogenic model. In rat astrocytes, DJ-1 overexpression protected against rotenone-induced neurotoxicity^63^. Overexpression of DJ-1 in mouse astrocytes was shown to be protective against rotenone-induced mitochondrial dysfunction and ROS generation of co-cultured neurons^13^. Our results and these studies highlight the relevance of DJ-1 in astrocytes and their impact on surrounding neurons, again emphasizing the importance of astrocyte dysfunction in the pathogenesis of PD. Concordant with the results in the astrocytes, knockdown of DJ-1 in the GBM cell lines also decreased GBM cell growth and metabolism, increased ROS and the immune response.

Thus, the mechanism by which DJ-1 modulates these pathways of cell proliferation, metabolism and immune response seems to be the same in both diseases. The opposite phenotypic observations in our models of PD and GBM can therefore be attributed to initially inverse DJ-1 levels, which modulate metabolic activity and downstream phenotypes. However, further studies are needed to reveal the exact molecular mechanism by which DJ-1 modulates the observed phenotypes in order to pave the way towards an improved therapy of these yet incurable conditions.

## Authors contribution

PM conducted the main experiments and drafted the manuscript. FJ and MMDLM performed brain section stainings. MU performed ICC experiments. FM supported main experiments. PKM and KB provided microglia. AJR helped with Seahorse, AR and GC performed qPCR experiments, PA and JJ helped with image analysis. SD, GD, FG and CJ helped with metabolomics, GA provided lentiviral constructs, AE established GSH assay, MP and JO performed RNAseq analysis, AL helped with Co-IP Mass spec. RT provided DJ-1 brain sections, DB and MM helped with brain section stainings. TS and LS helped with RNAseq. ZH, JCS, JM, AG, MP, IB and RK advised, supervised. FM and IB performed biotin switch assays. PM, IB and RK conceived the study and wrote the manuscript. All authors reviewed the manuscript for intellectual content and approved the current version for posting on bioRxiv.

## Methods

### Cell lines

We used isogenic pairs with two different homozygous DJ-1 mutations - P158Δ in-frame deletion (DelP) and DelP gene-corrected (GC), and c.192G>C (C4 mut) and C4, respectively^21–23^.

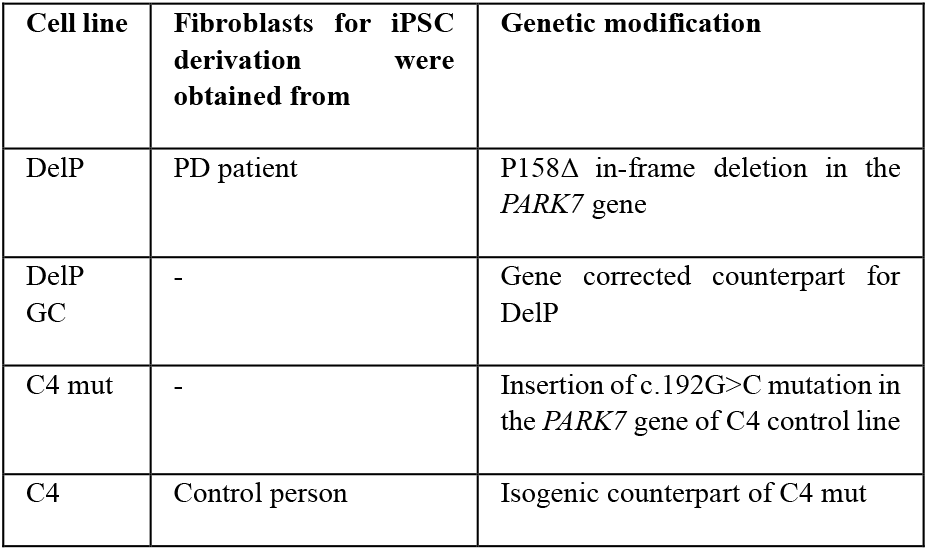

### Cell culture iPSC

Parkinson’s disease patient-derived iPSC of the DelP mutant and the isogenic control were generated as described by Mencke et al. 2022^22,23^. The C4 and C4 mut line (WT+DJ-1 mutant) were previously described^21^. All cells (iPSC, smNPC, hNSC, astrocytes and neurons) were cultivated in wells and flasks coated with Geltrex (Gibco™ A1413302). iPSCs were cultivated with DMEM/F12 (+Hepes) (Life/Tech – 31330038) supplemented with 10 % mTESR1 (STEMCELL Technologies SARL, 5850), 1 % insulin transferrin selenin (Life/Tech – 41400045), 1 % penicillin/streptomycin (Life/Tech – 15140-163), ascorbic acid 2PM (Sigma – A8960-5G) 64µg/mL, FGF-2 (Peprotech - 100-18B) 10 ng/mL, TGF-β1 (Peprotech - 100-21) 2 ng/mL and Heparin (Sigma – H3149-25KU) 100 ng/ml.

### smNPC

Differentiation of iPSC into smNPC was adopted from Reinhardt et al. 2013. On day one of differentiation, the iPSC medium was changed to iPSC medium without FGF-2 and mTESR1 plus 10 μM SB-431542 (Sigma - S4317-5mg), 1 μM dorsomorphin (Sigma – P5499-5mg), 3 μM CHIR 99021 (Axon – Axon1386) and 0.5 μM Purmorphamine (PMA) (Sigma-Aldrich – SML0868-25mg). On day 2, the medium was replaced by N2B27 medium: 50:50 DMEM/F12 w/o HEPES (Life/Tech – 21331046) and NeuroBasal medium (Life/Tech – 21103049) supplemented with 1:200 N2 supplement (Life/Tech 17502048), 1:100 B27 supplement lacking vitamin A (Life/Tech – 12587-010), 1 % penicillin/streptomycin (Life/Tech – 15140-163) and 1 % GlutaMAX Supplement (Life/Tech – 35050-061) plus 10 μM SB-431542, 1 μM dorsomorphin, 3 μM CHIR 99021 and 0.5 μM PMA. On day five, the medium was changed to N2B27 medium supplemented with 3 μM CHIR 99021, 0.5 μM PMA and 150 μM ascorbic acid (Sigma – A8960-5 g). Upon formation of neuroepithelial structures, the neuroepithelium was picked with a pipet tip, collected in a 1.5 ml tube, dissociated by pipetting and plated into 12-well in N2B27 medium supplemented with 3 μM CHIR 99021, 0.5 μM PMA and 150 μM ascorbic acid.

### Astrocytes

Astrocytes were generated from smNPC via hNSCs as described in Palm et al. 2015^64^. 2 days prior to hNSC differentiation, 400 k smNPCs were seeded into one 6-well of a 6-well plate per line. After 2 days, the medium was changed to smNPC medium with 20 ngng/ml FGF-2 (Peprotech - 100-18B). After 4 days, the cells were split with Accutase® (Sigma A6964) and the medium was changed to hNSC medium consisting of DMEM/F12 w/o HEPES (Life/Tech – 21331046) supplemented with N2 supplement (Life/Tech – 17502048), B27 supplement with vitamin A (Life Technologies Europe BV/Thermo Fisher Scientific 17504044), GlutaMAX Supplement (Life/Tech – 35050-061), penicillin/streptomycin (Life/Tech – 15140-163), 40 ng/ml EGF (Peprotech - AF-100-15-1mg), 40 ng/ml FGF-2 (Peprotech - 100-18B) and 1.5 ng/ml hLIF (Peprotech - AF-300-05). hNSCs were split with Accutase® when reaching 70-80 % of confluence. The astrocytic differentiation medium consisted of the basic cultivation medium DMEM/F12 w/o HEPES (Life/Tech – 21331046) supplemented with 1 % penicillin/streptomycin (Life/Tech – 15140-163), 1 % GlutaMAX Supplement (Life/Tech – 35050-061) and 1 % fetal bovine serum (Life/Tech 10270-106). 1 million hNSCs per T25 flask were plated 2 days prior to astrocyte differentiation. After 2 days, hNSC medium was changed to astrocyte medium. After 40 days, astrocytes were split to get rid of neurons that are dying during the differentiation and during the process of splitting. After 60 days, astrocytes were considered to be mature and all experiments were conducted around day 60.

### Glioblastoma cell culture

GBM cell lines LN229, U87 and U251 were kindly provided by Dr. Johannes Meiser from the LIH. Cells were cultured with DMEM 1X high glucose with glutamine (Thermo) and supplemented with 10 % FBS. Knockdown of DJ-1 was done using three different shRNAs (SigmaAldrich) with different efficiencies (#18, #19#, #20). GBM lines were seeded one day prior to transduction (500 k cells per well in 6-well plates for LN229 and 750 k cells per well in 6-well plates for U87 and U251). The next day, cells were transduced for 242424 hours in the presence of polybrene 8 μg/ml. 7 days later, cells were seeded for metabolite glucose tracing (100 k cells per well in 12-well plates), Western blottingblotting (1 million cells per well in 6-well plates) and RNA collection (500 k cells per well in 12-well plates).

In addition, a stable knockdown was generated for each GBM cell line using a mix of shRNA #18 and shRNA #20. Transduction was performed as described. The puromycin selection was started 24 hours post transduction for 4 days (U87: 1.5 μg/ml, U251: 1 μg/ml, LN229: 1.5 μg/ml).

### Microglia

Maintenance of iPSC lines (C4 healthy control and DJ-1 deficient C4 mut line) was done in mTeSR™ Plus medium (Stem Cell Technologies). To achieve microglia differentiation, a previously established protocol was implemented^55,56^ as described briefly in Badanjak et al. 2021^67^.

### Midbrain dopaminergic neuronal culture

Midbrain dopaminergic neurons were differentiated as described by Reinhardt et al. 2013. Cells were split on days 2 and 5 during differentiation and then cultivated until final seeding. Neurons were used for experiments from day 21 on.

### Generation of DJ-1 overexpression astrocytes

Wildtype iPSC were stably transduced with a GFP-containing DJ-1 overexpression vector. Cells were sorted with a FACS Aria sorter for GFP to obtain around 90 % GFP+ iPSC (Suppl. Figure 6A-C). iPSC were differentiated into smNPCs as described by Reinhardt and colleagues. smNPCs were sorted again for GFP to obtain 100 % GFP+ DJ-1 overexpressing cells (Suppl. Figure 6D). DJ-1 overexpression was confirmed by qPCR and Western blotting (Suppl. Figure 6E). Wildtype and DJ-1 overexpression astrocytes were differentiated into astrocytes and cultured as described by Palm and colleagues^66^ (Suppl. Figure 7). Characterization of astrocytes was performed by FACS. Wildtype iPSC-derived midbrain dopaminergic neurons (differentiated according to Reinhardt and colleagues) were used as negative control.

### Knockdown of DJ-1 in GBM cell lines

To assess the phenotypic effect of DJ-1 downregulation in the GBM cells, we used lentiviral constructs expressing three different shRNAs (Suppl. Fig. 10B-C). shRNA #20 had the strongest knockdown efficiency in all three lines and reduced the DJ-1 mRNA levels by around 80 % compared to shCtrl and the DJ-1 protein levels to around 50-60 % compared to shCtrl (Suppl. Fig. 10B-C).

### Growth assay

Astrocytes were seeded at a density of 100.000 cells per well in 12-well plates in duplicates and GBM cells were seeded at a density of 200.000 cells per well in 6-well plates in duplicates. Cells were counted at time points 0, 24, 48, 72 and 96 hours.

### Imaging

Antibodies used for imaging can be found in supplementary Table 1.

### Immunocytochemistry

iPSC, smNPC and hNSC were plated onto coverslips in 24-well plates (50.000 cells per well) and fixed with 4 % PFA prior to staining. Cells were stained for the typical markers listed in Table 1 using standard immunocytochemistry techniques. Images were acquired using a Zeiss spinning disk confocal microscope. Astrocytes, neurons and microglia were plated in CellCarrier-384 ultra Microplates (Perkin Elmer 6057300) (10.000 cells per well for neurons and astrocytes, 25.000 cells per well for microglia). All cells were fixed with 4 % PFA for 15 minutes. Cells were stained for the markers listed in Table 1 using standard immunocytochemistry methods. Images were taken using a Yokogawa cell voyager microscope. 16 images were taken per well in 384-well plates. Image analysis was performed using Matlab.

### Immunohistochemical staining

Sections of frontal cortex were cut at 6 µm thickness from formalin fixed, paraffin embedded blocks and mounted onto glass slides. Sections were stained automatically by using two equivalent Dako Omnis Autostainers (Dako), including hematoxylin counterstaining. According to the manufacturer’s instructions, each staining was performed by using default IHC protocols from the Omnis instrument software. Once stained, the tissue sections were dehydrated by rinsing them with EtOH and coverslipped following routine procedures. Finally, they were analyzed by using a brightfield microscope.

### Characterization of astrocytes

Astrocytes and neurons were detached in single cell suspension using Accutase® and centrifuged at 300 g for 3 minutes. Cells were washed 3 times with PBS, at 700 g for 5 minutes and fixed with 4 % PFA for 15 min. Cells were washed 3 times with PBS, at 700 g for 5 minutes and split into FACS tubes for the different stainings, before being resuspended in Saponin buffer (0.05 % Saponin/1 % BSA/PBS). Cells were incubated for 20-30 minutes at 4 °C. For unstained and isotype controls, a few µl of each cell line was mixed. After 30 minutes, cells were diluted in PBS, pelleted at 700 g for 5 minutes and resuspended in 50 µl of the primary antibody solution. Primary antibodies were prepared 1:50 in Saponin buffer (50 µl per tube). Cells were stained for GFAP, FoxA2, TH, TUJ1 and S100β or Recombinant Rabbit IgG, monoclonal [EPR25A] Isotype Control (ab172730) and Mouse IgG2a, kappa monoclonal [MG2a-53] Isotype control (ab18415). No primary antibody was added to the unstained control. Cells were incubated for 30 minutes at 4 °C. Cells were washed 3 times with diluted FACS buffer, centrifuged at 700 g for 5 minutes (FACS Buffer: PBS + 5 % BSA + 0.1 % Sodium Azide (NaN3), Diluted FACS Buffer: 1:5 dilution of FACS buffer). Cells were resuspended in secondary antibody solution (Alexa Fluor 568 Goat α-Rabbit IgG (H+L), Alexa Fluor 647 Goat α-Mouse IgG (H+L), 1:100 in PBS/10 % BSA, 50 µL per tube) and incubated for 30 minutes at 4 °C, washed 2x with diluted FACS buffer, IL-1β (H6291 Sigma) working conc. 10 ng/ml, Stock: 10ug/ml in 0.1% human albumin in PBS, use 1:1000 from stock (10ul aliquot) 0.1% human albumin in PBS control, use 1:1000 from 0.1% stock untreated control centrifuged at 700 g for 5 minutes, resuspend in 250 µL – 350 µL PBS and analyzed with BD LSRFortessa flow cytometry analyzer. The mean fluorescence intensity was assessed on single cells by using FlowJo LLC software.

### Annexin V assay

Astrocytes were deprived from glutamine in the medium for 4 hours. After 4 hours, cells were detached with Accutase® and centrifuged at 300 g for 3 minutes. Cell pellets were resuspended in 300 μL of 1X Annexin V binding buffer (Annexin-binding buffer 5X concentrate, Thermo Fisher Scientific B.V.B.A. V13246). 100.000 cells in 100 μL were used per sample and 5 μL of Annexin V, Alexa Fluor® 568 conjugate (Life/Tech Europe BV/Thermo Fisher Scientific A13202) were added, and the samples were incubated in the dark for 15 minutes. After 15 minutes, 400 μL of 1X Annexin V binding buffer were added. For FACS analysis, 1 μL of DAPI were added per tube, and samples were analyzed with BD LSRFortessa flow cytometry analyzer. The mean fluorescence intensity was assessed on single cells by using FlowJo LLC software.

### T cell migration assay

T cell medium (for 500 ml: IL2 (, Bio-Techne, 202-IL-010), CD3/CD 28 T cell activator (Stemcell, 10991), 450 ml T cell media IMDM (Gibco, 21980-032), 50 ml heat-inactivated FBS (10 %) (Gibco, 10500-064), 5 ml Pen/Strep (1 %) (Gibco, 15140-122), 5 ml non-essential amino acids (NEAA) (1 %) (Gibco, 11140-035), 0.5 ml β-mercaptoethanol (50 μM) (Gibco, 21985023)). T cell medium was adjusted to 1 % for the assay to avoid the generation of an FBS gradient that can lead to T cell migration.

Buffy coats were retrieved from different donors (each donor was treated as one biological replicate) from Red Cross Luxembourg (ethical approval proof available with author). PBMCs were isolated using SepMate™ PBMC Isolation Tubes (50 ml) and Lymphoprep™ Density Gradient Medium following standard procedures described by STEMCELL Technologies (https://www.stemcell.com/products/brands/sepmate-pbmc-isolation.html). PBMCs were frozen down until T cell isolation. PBMCs were cultivated in RPMI with 10 % FBS and 1 % penicillin/streptomycin. 11 days prior to the migration assay, PBMCs were thawed. T cells were isolated the day after using the Pan T cell isolation kit from Miltenyi following the manufacturer’s instructions. Cells were characterized by FACS on the day of isolation (FACS staining for CD3, CD4, and CD8). T cells were activated on the next day prior to viral transduction using ImmunoCult™ Human CD3/CD28 T Cell Activator (Stemcell Technologies, Catalogue #10971) following the ‘manufacturer’s manufacturer’s protocol. The next day, T cells were transduced with shCtrl and shRNA for DJ-1 #18, #19, and #20. The supernatant was removed after 24 hours and 3 days later, 200.000 astrocytes were seeded into a geltrex-coated bottom well of a 12 mm Transwell® with 3.0 µm Pore Polycarbonate Membrane Insert plate (Corning, 3402). Cells were either seeded in normal astrocyte medium or astrocyte medium supplemented with 5 mM NAC. After 2 days, astrocytes were stimulated with

10 ng/ml IL-1β in either normal astrocyte medium, astrocyte medium containing 5 mM NAC or 4 mM glutamine (double amount). The next day, T cells were activated again using ImmunoCult™ Human CD3/CD28 T Cell Activator. The day after (T cells were already 10 days in culture), inserts were placed into the stimulated astrocytes and 100.000 T cells were added in T cell medium with 1% FBS only into each insert. One well was kept without astrocytes as passive migration control and one well contained only astrocyte medium with CCL5 as positive control. Cells were incubated for 4 hours. After 4 hours, the transwell filter was removed and the T cells present in the upper chamber were saved. The astrocyte medium in the lower chamber was centrifuged to retrieve migrated T cells. The medium was saved for ELISA to analyze cytokine release after 48 hours upon IL-1β stimulation. T cells were analyzed by FACS using the antibodies listed in Supplementary Table 1 and CountBright™ Absolute Counting Beads following the manufacturer’s instructions. Stopping gate was CD3.

### Interleukin 1 beta (IL-1β) treatment

Seed 200.000 astrocytes per 12well per condition. After 48hours, add the respective treatments (1 ml per 12well):

- IL-1β (H6291 Sigma) working conc. 10 ng/ml, Stock: 10ug/ml in 0.1% human albumin in PBS, use 1:1000 from stock (10ul aliquot)
- 0.1% human albumin in PBS control, use 1:1000 from 0.1% stock
- untreated control

After 48 hours of treatment, remove medium, add accutase and collect the cells after 5min. Spin 3min 500g, remove supernatant and store the pellets at -80°C until RNA extraction and subsequent cDNA synthesis and qPCR.

### RNA extraction, cDNA synthesis, qPCR

RNA extraction was performed using the RNeasy Kit fromfrom Qiagen according to manufacturer’s instructions. RNA concentration was assessed using a NanoDrop™ spectrophotometer. Subsequent cDNA synthesis was done with the High-Capacity cDNA Reverse Transcription Kit with Rnase Inhibitor (Applied Biosystems™) following the manufacturer’s instructions. qPCR was done with the hDMSPhigh-throughput platform at the LCSB using the Echo® Acoustic Liquid Handling droplet ejection system. qPCRs were run with a LightCycler® 480 machine (40 cycles per run). Standard curves for each primer were included to assess the efficiency of each primer and subsequently calculate the Pfaffl ratio.

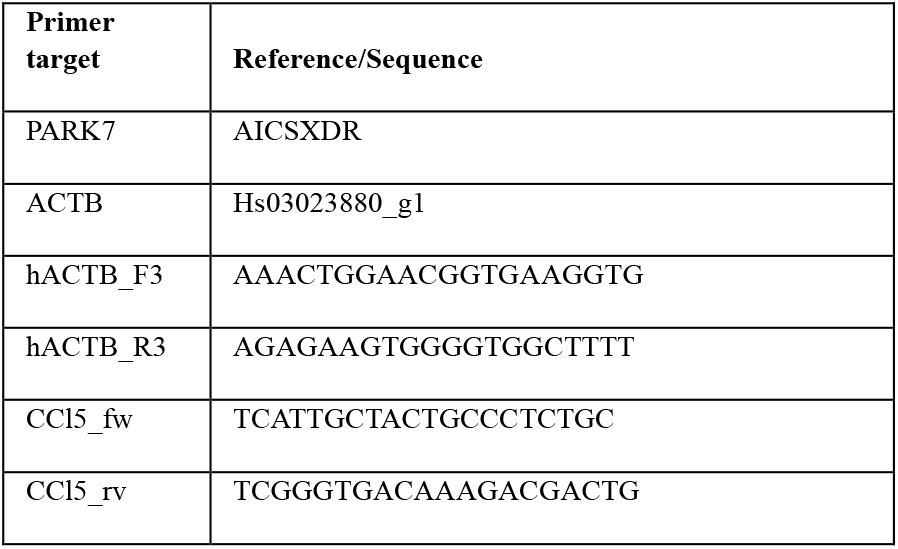

### RNA sequencing

RNA for RNA sequencing was extracted as described above (chapter: RNA extraction, cDNA synthesis, qPCR). RNA quality was assessed using the Agilent 2100 Bioanalyzer, RNA integrity (RIN) values were >8. Libraries were prepared using the TruSeq Stranded mRNA library prep kit and sequenced on a NextSeq2000. For GBM samples paired reads of 51 bp length were generated, for astrocyte samples single reads of 75 bp length were generated.

Data was processed using an in-house snakemake^68^ workflow available as a git repository https://git-r3lab.uni.lu/aurelien.ginolhac/snakemake-rna-seq (release v0.2.3, and singularity image v0.4). Raw read quality was assessed by FastQC (v0.11.9)^69^. Adapters are removed using AdapterRemoval (v2.3.1)^70^, with a minimum length of the remaining reads set to 35 bp. Reads were mapped to hg38 (GRCh38.p13) using STAR (v.2.7.4a)^571^, featureCounts from the R package Rsubread (2.2.2)^582^ was used to count reads. All counts >10 were used for differential gene expression analysis using the R package DESeq2 (v1.28.1)^73^. Normalization in DEseq2 was done using apeglm (v.1.10.0)^594^. All FPKM were calculated using DESeq2 package. Pathway analysis on DEGs with false discovery rate < 0.1 and a minimum log2-fold change cut-off of +/-0.5 was performed using Ingenuity Pathway Analysis tool from Qiagen, Content version: 60467501 (release date: 2020-11-19).

### Biotin Switch Assay

Biotin switch assay was conducted following a modified version of the protocol published by Butturini and colleagues^75^. In brief, lysates were treated with 50 mM N-Ethylmaleimide (NEM) to stably alkylate free thiol groups. Then, free thiol groups that were not alkylated by NEM were reduced by treatment with 60 mM Dithiothreitol (DTT). To remove free NEM and DTT, samples were cleaned using Amicon Utra Spin Desalting Columns (Millipore) following the manufacturer’s instructions. Next, free thiol groups were oxidized with 1 mM Diamide before incubation with 1 mM biotinylated glutathione (BioGEE) followed by treatment in Amicon Utra Spin Desalting Column (Millipore) to remove cellular GSH and oxidants in excess. Samples were suspended in cold RIPA Buffer supplemented with protease inhibitors prior to protein quantification. 5-10 µg protein of the lysate was saved for Western blot analysis of total DLAT and DLST content and staining of the housekeeping protein Vinculin. Of the remaining lysate 250 – 400 µg of proteins per sample were incubated for 1 h at 4° C with 100 uL of Slurry agarose beads under gentle rotation to reduce non-specific binding and background. For each set of parallel differentiations (wt, mut, mut Nac and mut hGln) the same amount of protein was used in the assay. Streptavidin-agarose beads were used to pull-down biotinylated proteins from the cleared lysate for 2 h at 4° C under gentlre rotation. Streptavidin pull-down samples were separated on a 5–10% SDS–polyacrylamide gel (SDS-PAGE) in running buffer following the manufacturer’s instructions. Western Blot was conducted following standard procedures as described below using antibodies against DLAT and DLST (Supplementary Table 1).

### Western Blotting

Cells were lysed with 200 µL lysis buffer (1 % SDS + protease inhibitor cocktail tablet Roche) per well in 6-well plates and the lysate was collected by scraping. The lysate was transferred into 1.5 ml tubes and boiled for 5 minutes at 95 °C, centrifuged briefly and stored at -80 °C. Samples were sonicated after thawing. Protein quantification was performed using the Pierce BCA assay. Invitrogen™ NuPAGE™ 4 to 12 %, Bis-Tris, 1.0 mm, Mini Protein Gels with 12-wells were used for all blots. Prior to loading, samples were diluted with loading buffer (NuPAGE™ LDS Sample Buffer (4X) with RA 10x) and boiled for 5 minutes at 95 °C. One ml of anti-oxidant for 400 ml of running buffer (NuPAGE™ MES SDS Running Buffer (20X)) were used. As ladder, PageRuler Plus Prestained Protein Ladder was used. After the run, wet transfer was performed following standard protocols (transfer buffer 1:5, 99 % Ethanol, 1X Tris glycine in MilliQ water). The transfer was run for 75 minutes at 100 V. Ponceau red was always used after the transfer. The membrane was rinsed with TBS-T and blocked with 5 % milk in TBS-T for 1 hour prior to incubation of the primary antibody overnight at 4 °C with rotation. The next day, the membrane was washed 3 times for 10 minutes in TBS-T with shaking and incubated with the secondary antibody for 1 hour at room temperature with shaking. After 1 hour, the membrane was washed 3 times for 10 minutes in TBS-T with shaking before revealing the gel using an Odyssey® Imager after incubating the membrane for 30 seconds with ECL solution: 50 % Peroxide solution + 50 % Luminol solution.

### Metabolic carbon contribution analysis using gas chromatography – mass spectrometry (GC-MS) and Liquid chromatography – mass spectrometry (LC-MS)

#### Intracellular metabolite extraction

Stable isotope-assisted metabolomics analyses were conducted using [U-13C6]Glucose or [U-13C5]Glutamine tracers. Cells were incubated with the tracing medium for 48 hours to reach an isotopic steady-state (200.000 cells per well in 12-well plates, with technical triplicates). After 48 hours, intracellular metabolite extraction was performed at 4 °C. First, the medium was collected for further extracellular metabolite analysis. For [U-13C6] Glucose labelling, measured using GC-MS, cells were washed once with 1 mL 0.9 % NaCl solution. Then, 200 μL of methanol were added (containing 5 µg/mL Tridecanoid-*D25* acid as internal standard), followed by the addition of 80 μL of MilliQ water (4 °C) (containing 1 µg/mL Pentanedioic-*D6* acid as internal standard). The plates were gently shaken for 10 minutes at 4 °C before the mixture was transferred into a new 1.5 mL-reaction tube containing 100 uL of Chloroform. The reaction tubes were shaken for 5 minutes at 4 °C and full speed in an Eppendorf Thermomixer. For phase separation, 100μL of Chloroform and 100 uL of water were added. Afterwards, the reaction tubes were vigorously vortexed for 10 seconds and centrifuged for 5 minutes at 4°C and full speed. 125 μL of the upper polar phase were transferred into a GC vial with micro insert. The samples were evaporated in a centrifugal vacuum concentrator at -4 °C, capped and stored at -80 °C until GC-MS measurement.

For LC-MS analyses, cells were washed once with 1 mL 0.9 % NaCl solution. Then, 250 μL of an extraction fluid (4:1, Methanol/H2O mixture) were added to each well. The following internal standards were added to the water fraction of the extraction fluid: [UL-^13^C]Ribitol (*c* = 2 μg/mL), Pentanedioic-*D6* acid (*c* = 2 μg/mL), Tridecanoic-*D25* acid (*c* = 10 μg/mL), 6-Chloropurine riboside (*c* = 10 μg/mL), 4-Chloro-DL-phenylalanine (*c* = 10 μg/mL), Nε-Trifluoroacetyl-L-lysine (*c* = 10 μg/mL), Thionicotinamide adenine dinucleotide (*c* = 10 μg/mL). Then, 30 μL of MilliQ water (4 °C) were added per well and the procedure was continued like described above.

#### Extracellular metabolite extraction

Medium was filtered using a Phenex Regenerated Cellulose (RC) syringe filter prior to freezing to remove any cells or debris. 20 μL of the sample (spent medium, fresh medium and calibrants) were added to 180 μL ice-cold extraction fluid (5:1, Methanol/H2O mixture). Two internal standards, consisting of [U-13C]Ribitol (*c* =final concentration: 50 µg/mL) and Pentanedioic-*D6* acid (final concentration: 20 µg/mL), were added to the water fraction of the extraction fluid. Then, samples were vortexed for 10 seconds and incubated for 15 minutes at 4 °C at maximum speed in an Eppendorf Thermomixer, followed by centrifugation for 5 minutes at 4 °C and full speed. 50 μL of the supernatant were transferred into a GC vial with micro insert. The samples were evaporated in a centrifugal vacuum concentrator at-4 °C, capped and stored at -80 °C until GC-MS measurement.

#### Metabolomics data acquisition and data analysis

Sample measurements were performed at the LCSB Metabolomics Platform using a set of GC-MS (incl. derivatization) and LC-MS methods to analyze intracellular^60,61^ and extracellular^60^ metabolites. Raw mass spectrometric data was manually curated and normalized to internal standards (quantification) or corrected for natural isotopic abundance (determination of MIDs) before MS data was exported.

Explanation of metabolite tracing illustrated in supplementary Figure 14.

#### Intracellular

R was used to analyze the MS data. Details on all packages used can be found in the code deposited at https://gitlab.lcsb.uni.lu/TNG/papers/PD_GBM_publication. Raw data (mass isotopomer distribution for each metabolite) were loaded as excel files and formatted according to tidy data guidelines^62^. The means of the three independent replicates per biological replicate were calculated for each metabolite before calculating the mean of the biological replicates. Data were plotted in R using ggplot. Means and single values for each metabolite (isotopologue fractions) per sample were exported as .csv files and carbon contribution was calculated from isotopologue fraction for each metabolite as follows: for example for a 5 carbon molecule: (M1*1 + M2*2 + M3*3 + M4*4 + M5*5) /5. Final graphs were generated using GraphPad Prism. Full MIDs can be found deposited at https://gitlab.lcsb.uni.lu/TNG/papers/PD_GBM_publication and as supplementary Figures 12 and 13.

#### Extracellular

After loading the raw data (internal standard normalized peak areas), data were formatted according to tidy data guidelines. Outliers in standards were removed using interquartile range. To determine the concentration of each metabolite in the cell culture tracing medium and the fresh tracing medium, the calibration curve (measured in technical triplicates) was used to apply linear regression and calculate the unknown concentration for each metabolite. The concentration of each metabolite in Mol was subtracted by the concentration in Mol for each metabolite measured in the fresh tracing medium to obtain a delta corresponding to uptake and release of the respective metabolite in Mol. The means of the three independent replicates per biological replicate were calculated for each metabolite in Mol before calculating the mean of the biological replicates. Data were again plotted in R using ggplot. Means and single values for each metabolite per sample were also exported as .csv files and final graphs were generated using GraphPad Prism.

#### Seahorse

Oxygen consumption rate (OCR) and Extracellular Acidification Rate (ECAR) as well as the glycolytic stress test were measured in whole cells using the Seahorse Xfe96 Cell Metabolism Analyzer (Agilent). The concentrations of mitochondrial toxins and compounds used were optimized according to the manufacturer’s recommendations. Final concentrations are listed in Table 2.

**Table 2.**

| Compound | Reference | Company | Molecular weight | Final concentration for assay |
| --- | --- | --- | --- | --- |
| *Oligomycin* | Lot APN14317-13, ab141829 | Abcam | 791.062 | 10 µM for OCR |
| *Oligomycin* | Lot APN14317-13, ab141829 | Abcam | 791.062 | 100 µM for glycolytic stress test |
| *FCCP* | C2920-10MG | Sigma | 254.1681 | 2 µM |
| *Rotenone* | R8875-1G | Sigma | 394.42 | 5 µM |
| *Antimycin A* | A86474 | Sigma | 532 | 5 µM |
| *Glucose Monohydrate* | 6887 | Carl Roth | 180.156 | 80 mM |
| *2-DG* | D8375-5G | Sigma | 164.16 | 500 mM |

Compound aliquots were dissolved in Seahorse Base medium on the day of the experiment. To prepare the Seahorse Base medium, 990 mL of ddH2O were autoclaved and 1 vial of DMEM Basal Powder (Sigma D5030) was added, the solution was sterile filtered, and the pH was adjusted to 7.4 ± 0.05 at 37 °C (pH was checked for every experiment). Seahorse assay medium for the two different assays was prepared according to the manufacturer’s instructions.

Laminin (Sigma, L2020; 25 µL aliquot) was added to 2.55 mL of PBS before 25 µL of diluted laminin were added into each well of a Seahorse Xfe96 well plate. The plate was incubated overnight at 37 °C. Astrocytes were plated in the Seahorse Xfe96 well plate 24 hours prior to measuring at a density of 80.000 cells per well. Perimeter wells were not used due to evaporation. After seeding, the plate was left at room temperature for 1 hour to prevent edge-effects. After 1 hour, all unused wells were filled with astrocyte media only and incubated overnight at 37 °C + 5 % CO2. The cartridge was hydrated using 200 µL of XF-calibrant solution and incubated overnight at 37 °C in a non-CO2 incubator. 100 mL of each of the prepared Seahorse assay media were aliquoted, the pH was adjusted to 7.4 ± 0.05 at 37 °C and the media were incubated in a non-CO2 incubator overnight. The media of the cells was changed approximately 1 hour before starting the experiment by removing 60 µL of culture media (leaving 20 µL) from each well, rinsing 2x with 200 µL of Seahorse assay media and eventually adding 155 µL of assay media to each well for a final volume of 175 µL/well. The plate was then incubated in non-CO2 incubator for at least an hour so that the cells were allowed to equilibrate to the assay media. The compounds were loaded into the cartridge on the Seahorse utility plate using the loading guide plates while the cells are incubating in the non-CO2 incubator. After the loading of the compounds, the plate was incubated for 30 minutes in a non-CO2 incubator before the measurement was started. Normalization was performed using the CyQUANT® kit. After the Seahorse assay, the assay media was removed from the wells by inverting and blotting the surplus onto paper towels and the Seahorse cell culture plates were stored at -80 °C until the CyQUANT® assay was performed. The CyQUANT® GR stock solution in DMSO was brought to room temperature. The Seahorse cell culture plates were also equilibrated at room temperature. The 20x concentrated cell-lysis buffer stock solution (Component B) was diluted 20-fold in distilled water. For each well, 200 μL were required. The CyQUANT® GR stock solution (Component A) was diluted 400-fold into the 1X cell-lysis buffer. 200 μL of the CyQUANT® GR dye/cell-lysis buffer were added to each well. The plates were incubated 2–5 minutes at room temperature, protected from light. The samples were transferred into a 96-well plate before measuring the sample fluorescence using a fluorescence microplate reader with filters appropriate for ∼480 nm excitation and ∼520 nm emission maxima.

#### ROS assay

5000 GBM cells and 5000 astrocytes were seeded into a CellCarrier-384 ultra Microplate (Perkin Elmer 6057300). For NAC treated cells, cells were seeded with medium containing 5 mM NAC. Two days later, the medium was changed and for the hGln condition (GlutaMAX Supplement (Life/Tech – 35050-061)), medium with hGln (4 mM for astrocytes medium, 8 mM for GBM cell medium) was added. Two days after that, cells were starved for 4 hours without FBS. To analyze ROS, the ROS-ID® Total ROS/Superoxide detection kit - ENZ-51010 was used. The assay was conducted following the manufacturer’s instructions (except no pretreatment with NAC for 30 minutes was necessary as respective wells (NAC treated) had NAC for 9696 hours). After the assay, plate contents were emptied and 20 μL of RIPA buffer were added per well. The plate was stored for 30 minutes at -80 °C. After 30 minutes, the plate was retrieved from the freezer and the lysis buffer was thawed. Once the buffer was thawed, Pierce BCA was performed according to the manufacturer’s protocol (30 μL of BCA working solution per well).

#### GSH/GSSG measurement

Cells were seeded in a 96-well plate with different media (20k cells per well). Cells were cultivated for 48 hours. Two days later, cells were detached, counted and 3000 cells per 384-well for each condition were centrifuged. The GSH and GSSG measurement was performed using the GSH GSH/GSSG-Glo™ Assay from Promega following the manufacturer’s instructions in 384-well format.

## Data analysis

Each GBM cell line was treated as biological replicate so for each line, experiments were run in technical replicates and statistics were calculated using the mean of all the data. Metabolomics data were analyzed using R. Images were analyzed using the Zeiss Zen program and Yokogawa images were analyzed using Matlab and R. All original code has been deposited at https://gitlab.lcsb.uni.lu/TNG/papers/PD_GBM_publication. FACS data were analyzed using FlowJo LLC software. Final graphs were made using GraphPad Prism. All experiments have sample sizes equal to or higher than 3. Exclusion criteria were biological and technical variation. Statistical method of comparison was paired t-test for the two isogenic pairs. All error bars show the standard error of the mean.

## Materials availability statement

All cell lines are available upon request. More information on use restrictions and on how to submit the request to obtain the cell lines is available at DOI: 10.17881/0m4p-ht15.

## Data and code availability

The raw RNA sequencing data for this manuscript are not publicly available due to its sensitive nature. The data are available upon request. All derived data, data behind figures and supplementary material has been deposited at https://data.mendeley.com/datasets/xv5gt4hpjd/draft?a=f39d6035-c276-4c2b-90c6-e8f21b4cffdd under CC-BY license and it will be publicly available at DOI: 10.17632/xv5gt4hpjd.

More information on availability of data and how to submit the request to access the sensitive dataset is available at DOI: 10.17881/0m4p-ht15.

Code is deposited at https://gitlab.lcsb.uni.lu/TNG/papers/PD_GBM_publication.

Any additional information required to reanalyze the data reported in this paper is available from the lead contact upon request.

## Acknowledgements

We would like to thank the patient for providing fibroblasts for the generation of the described cell line DelP and DelP GC. We would like to thank the LCSB Metabolomics Platform for providing technical and analytical support. We acknowledge Portuguese Brain Bank for tissue samples supply. We acknowledge DMSP for the help with qPCRs, we acknowledge the former heads of DMSP Yong-Jun Kwon and Meritxell B. Cutrona. The graphical abstract was created with BioRender.com.

## Funding

This current work was supported by the Fonds National de Recherche (FNR) within the PEARL Excellence Programme [FNR/P13/6682797] to RK, and the MiRisk project [C17/BM/11676395], by the Jean Think Foundation Luxembourg, FNR-ATTRACT program (A18/BM/11809970) to JM, FNR [AFR PhD 12447024] to PM. NCER-PD supported work of RK. MAMA-Syn funding supported JO. FNR (PRIDE17/11823097/MicrOH) and FNR-CORE (C21/BM/15796788) supported AE and DB.

## Competing interests

The authors report no competing interests.

## Notes

https://gitlab.lcsb.uni.lu/TNG/papers/PD_GBM_publication

### Competing Interest Statement

The authors have declared no competing interest.

## References

1. Booth, H. D. E., Hirst, W. D. & Wade-Martins, R. The Role of Astrocyte Dysfunction in Parkinson’s Disease Pathogenesis. Trends Neurosci 40, 358–370 (2017).

2. Miyazaki, I. & Asanuma, M. Neuron-Astrocyte Interactions in Parkinson’s Disease. (2020) doi:10.3390/cells9122623.

3. Vergara, R. C. et al. The Energy Homeostasis Principle: Neuronal Energy Regulation Drives Local Network Dynamics Generating Behavior. Front Comput Neurosci 13, 1–18 (2019).

4. Turner, D. A. & Adamson, D. C. Neuronal-astrocyte metabolic interactions: Understanding the transition into abnormal astrocytoma metabolism. J Neuropathol Exp Neurol 70, 167–176 (2011).

5. Kam, T. I., Hinkle, J. T., Dawson, T. M. & Dawson, V. L. Microglia and astrocyte dysfunction in parkinson’s disease. Neurobiol Dis 144, 105028 (2020).

6. Wang, B. et al. Effect of DJ-1 overexpression on the proliferation, apoptosis, invasion and migration of laryngeal squamous cell carcinoma SNU-46 cells through PI3K/AKT/mTOR. Oncol Rep 32, 1108–1116 (2014).

7. Smajic, S. et al. Single-cell sequencing of human midbrain reveals glial activation and a Parkinson-specific neuronal state. Brain 145, 964–978 (2022).

8. Ariga, H. Common mechanisms of onset of cancer and neurodegenerative diseases. Biol Pharm Bull 38, 795–808 (2015).

9. Bonifati, V. et al. Mutations in the DJ-1 gene associated with autosomal recessive early-onset parkinsonism. Science (1979) 299, 256–259 (2003).

10. Canet-Avilés, R. M. et al. The Parkinson’s disease DJ-1 is neuroprotective due to cysteine-sulfinic acid-driven mitochondrial localization. Proc Natl Acad Sci U S A 101, 9103– 9108 (2004).

11. Bandopadhyay, R. et al. The expression of DJ-1 (PARK7) in normal human CNS and idiopathic Parkinson’s disease. Brain 127, 420–430 (2004).

12. Rizzu, P. et al. DJ-1 Colocalizes with Tau Inclusions: A Link between Parkinsonism and Dementia. Ann Neurol 55, 113–118 (2004).

13. Mullett, S. J., Di Maio, R., Greenamyre, J. T. & Hinkle, D. A. DJ-1 Expression Modulates Astrocyte-Mediated Protection Against Neuronal Oxidative Stress. Journal of Molecular Neuroscience 49, 507–511 (2013).

14. Mencke, P. et al. The Role of DJ-1 in Cellular Metabolism and Pathophysiological Implications for Parkinson’s Disease. Cells 10, 1–18 (2021).

15. Bélanger, M., Allaman, I. & Magistretti, P. J. Brain energy metabolism: Focus on Astrocyte-neuron metabolic cooperation. Cell Metab 14, 724–738 (2011).

16. Wang, C. et al. The positive correlation between DJ-1 and β-catenin expression shows prognostic value for patients with glioma. Neuropathology 33, 628–636 (2013).

17. Goffart, N., Kroonen, J. & Rogister, B. Glioblastoma-initiating cells: Relationship with neural stem cells and the micro-environment. Cancers (Basel) 5, 1049–1071 (2013).

18. Jiang, Y. & Uhrbom, L. On the origin of glioma. Ups J Med Sci 117, 113–121 (2012).

19. Taipa, R. et al. DJ-1 linked parkinsonism (PARK7) is associated with Lewy body pathology. Brain 139, 1680–1687 (2016).

20. Wilhelmsson, U. et al. Redefining the concept of reactive astrocytes as cells that remain within their unique domains upon reaction to injury. Proc Natl Acad Sci U S A 103, 17513–17518 (2006).

21. Boussaad, I. et al. A patient-based model of RNA mis-splicing uncovers treatment targets in Parkinson’s disease. Sci Transl Med 12, (2020).

22. Mencke, P. et al. Generation and characterization of a genetic Parkinson’s disease-patient derived iPSC line DJ-1-delP (LCSBi008-A). Stem Cell Res 62, 1–5 (2022).

23. Mencke, P. et al. Generation of isogenic control DJ-1-delP GC13 for the genetic Parkinson’s disease-patient derived iPSC line DJ-1-delP (LCSBi008-A-1). Stem Cell Res 62, 102815 (2022).

24. Qiagen, C. version: 60467501 (release date: 2020-11-19). IPA.

25. Turner, D. A. & Adamson, D. C. Neuronal-astrocyte metabolic interactions: Understanding the transition into abnormal astrocytoma metabolism. J Neuropathol Exp Neurol 70, 167–176 (2011).

26. Meiser, J. et al. Loss of DJ-impairs antioxidant response by altered glutamine and serine metabolism. Neurobiol Dis 89, 112– 125 (2016).

27. Zhang, L. et al. Role of DJ-1 in Immune and Inflammatory Diseases. Front Immunol 11, 1–10 (2020).

28. Eltoweissy, M., Dihazi, G. H., Müller, G. A., Asif, A. R. & Dihazi, H. Protein DJ-1 and its anti-oxidative stress function play an important role in renal cell mediated response to profibrotic agents. Mol Biosyst 12, 1842–1859 (2016).

29. Xu, X. M. & Møller, S. G. ROS removal by DJ-1: Arabidopsis as a new model to understand Parkinson disease. Plant Signal Behav 5, 1034–1036 (2010).

30. Waak, J. et al. Regulation of astrocyte inflammatory responses by the Parkinson’s disease-associated gene DJ-1. FASEB Journal 23, 2478–2489 (2009).

31. Clements, C. M., McNally, R. S., Conti, B. J., Mak, T. W. & Ting, J. P. Y. DJ-1, a cancer- and Parkinson’s disease-associated protein, stabilizes the antioxidant transcriptional master regulator Nrf2. Proc Natl Acad Sci U S A 103, 15091–15096 (2006).

32. Ariga, H. et al. Neuroprotective function of dj-1 in Parkinson’s disease. Oxid Med Cell Longev 2013, (2013).

33. Irrcher, I. et al. Loss of the Parkinson’s disease-linked gene DJ-1 perturbs mitochondrial dynamics. Hum Mol Genet 19, 3734– 3746 (2010).

34. Krebiehl, G. et al. Reduced basal autophagy and impaired mitochondrial dynamics due to loss of Parkinson’s disease-associated protein DJ-1. PLoS One 5, (2010).

35. Tavassolifar, M. J., Vodjgani, M., Salehi, Z. & Izad, M. The Influence of Reactive Oxygen Species in the Immune System and Pathogenesis of Multiple Sclerosis. Autoimmune Dis 2020, (2020).

36. Moreira Franco, Y. E. et al. Glutaminolysis dynamics during astrocytoma progression correlates with tumor aggressiveness. Cancer Metab 9, 1–15 (2021).

37. Obara-Michlewska, M. & Szeliga, M. Targeting glutamine addiction in meningioma. Neuro Oncol 24, 569–570 (2020).

38. Zong, H., Verhaak, R. G. W. & Canolk, P. The cellular origin for malignant glioma and prospects for clinical advancements. Expert Rev Mol Diagn 12, 383–394 (2012).

39. Salemi, M. et al. Examples of Inverse Comorbidity between Cancer and Neurodegenerative Diseases: A Possible Role for Noncoding RNA. Cells 11, 1930 (2022).

40. Mogi, M. Interleukin-lfl, interleukin-6, epidermal growth factor and transforming growth factor-are elevated in the brain from parkinsonian patients. 180, 147–150 (1994).

41. Sonninen, T. M. et al. Metabolic alterations in Parkinson’s disease astrocytes. Sci Rep 10, (2020).

42. Kim, J. hyeon et al. DJ-1 facilitates the interaction between STAT1 and its phosphatase, SHP-1, in brain microglia and astrocytes: A novel anti-inflammatory function of DJ-1. Neurobiol Dis 60, 1–10 (2013).

43. Floss, T., Angelis, D., Kahle, P. J. & Wurst, W. DJ-1-deficient mice show less TH-positive neurons in the ventral tegmental area and exhibit non-motoric. 305–317 (2010) doi:10.1111/j.1601-183X.2009.00559.x.

44. Chen, X., Song, M., Zhang, B. & Zhang, Y. Reactive Oxygen Species Regulate T Cell Immune Response in the Tumor Microenvironment. Oxid Med Cell Longev 2016, 11–16 (2016).

45. Yang, Z., Min, Z. & Yu, B. Reactive oxygen species and immune regulation. Int Rev Immunol 39, 292–298 (2020).

46. Naik, E. & Dixit, V. M. Mitochondrial reactive oxygen species drive proinflammatory cytokine production. Journal of Experimental Medicine 208, 417–420 (2011).

47. Ho, M., Yen, C., Hsieh, T., Kao, T. & Chiu, J. Redox Biology CCL5 via GPX1 activation protects hippocampal memory function after mild traumatic brain injury. Redox Biol 46, 102067 (2021).

48. Pittaluga, A. CCL5-glutamate cross-talk in astrocyte-neuron communication in multiple sclerosis. Front Immunol 8, 1–13 (2017).

49. Brochard, V. et al. Infiltration of CD4+ lymphocytes into the brain contributes to neurodegeneration in a mouse model of Parkinson disease. Journal of Clinical Investigation 119, 182– 192 (2009).

50. Singh, Y. et al. Differential effect of DJ-1 / PARK7 on development of natural and induced regulatory T cells. Nature Publishing Group 1–14 doi:10.1038/srep17723.

51. Belikov, A. V., Schraven, B. & Simeoni, L. T cells and reactive oxygen species. J Biomed Sci 22, 1–11 (2015).

52. Stanke, K. M., Wilson, C. & Kidambi, S. High Expression of Glycolytic Genes in Clinical Glioblastoma Patients Correlates With Lower Survival. Front Mol Biosci 8, 1–12 (2021).

53. Wang, H. et al. Different T-cell subsets in glioblastoma multiforme and targeted immunotherapy. Cancer Lett 496, 134– 143 (2021).

54. Mohan, A. A. et al. Targeting Immunometabolism in Glioblastoma. Front Oncol 11, 1–16 (2021).

55. Haenseler, W. et al. A Highly Efficient Human Pluripotent Stem Cell Microglia Model Displays a Neuronal-Co-culture-Specific Expression Profile and Inflammatory Response. Stem Cell Reports 8, 1727–1742 (2017).

56. van Wilgenburg, B., Browne, C., Vowles, J. & Cowley, S. A. Efficient, Long Term Production of Monocyte-Derived Macrophages from Human Pluripotent Stem Cells under Partly-Defined and Fully-Defined Conditions. PLoS One 8, (2013).

57. Dobin, A. et al. STAR: Ultrafast universal RNA-seq aligner. Bioinformatics 29, 15–21 (2013).

58. Liao, Y., Smyth, G. K. & Shi, W. The R package Rsubread is easier, faster, cheaper and better for alignment and quantification of RNA sequencing reads. Nucleic Acids Res 47, (2019).

59. Zhu, A., Ibrahim, J. G. & Love, M. I. Heavy-Tailed prior distributions for sequence count data: Removing the noise and preserving large differences. Bioinformatics 35, 2084–2092 (2019).

60. Becker, B. et al. Serine metabolism is crucial for cGAS-STING signaling and viral defense control in the gut. iScience 27, (2024).

61. Heins-Marroquin, U. et al. CLN3 deficiency leads to neurological and metabolic perturbations during early development. Life Sci Alliance 7, (2024).

62. Elliott, A. C., Hynan, L. S., Reisch, J. S. & Smith, J. P. Preparing data for analysis using Microsoft Excel. Journal of Investigative Medicine 54, 334–341 (2006).

