## Supplementary Material for "DJ-1 mediates regulation of metabolism and immune response in Parkinson’s disease astrocytes and Glioblastoma cells"

*Supplementary Table 1: Antibody list*

| Cell type to be stained | Antibody | Species | Company | Reference | Dilution | 2ndary antibody all 1:1000 for ICC, for FACS 1:100 |
|---|---|---|---|---|---|---|
| iPSC | Anti Nanog | rabbit | abcam | ab21624 | dilution 1:1000 | Alexa Fluor 568 Goat α-Rabbit IgG (H+L) A11036 |
| iPSC | Anti Oct3/4 | mouse | santa cruz | sc-5279 | dilution 1:1000 | Alexa Fluor 647 Goat α-Mouse IgG (H+L) A21236 |
| iPSC, smNPC | Anti SOX2 (Y-17) | goat | santa cruz | sc-17320 | dilution 1:250 | Alexa Fluor 647 Donkey α-Goat IgG (H+L) A21447 |
| smNPC, hNSC | Anti Nestin | mouse | R&D Systems | MAB1259 | dilution 1:1000 | Alexa Fluor 647 Goat α-Mouse IgG (H+L) A21236 |
| smNPC, hNSC | Anti Musashi | rabbit | abcam | ab21628 | dilution 1:250 | Alexa Fluor 568 Goat α-Rabbit IgG (H+L) A11036 |
| hNSC | Anti SOX1 | goat | R&D Systems | AF3369 | dilution 1:250 | Alexa Fluor 647 Donkey α-Goat IgG (H+L) A21447 |
| Astrocytes and neurons | Anti ID3 | mouse | abcam | ab236505 | dilution 1:1000 | Alexa Fluor 647 Goat α-Mouse IgG (H+L) A21236 |
| Astrocytes and neurons | Anti NFIA | rabbit | abcam | ab228897 | dilution 1:1000 | Alexa Fluor 568 Goat α-Rabbit IgG (H+L) A11036 |
| Astrocytes and neurons | Anti EZRIN | rabbit | abcam | ab40839 | dilution 1:500 | Alexa Fluor 568 Goat α-Rabbit IgG (H+L) A11036 |
| Astrocytes and neurons | Anti S100b | rabbit | abcam | ab868 | dilution 1:500 | Alexa Fluor 568 Goat α-Rabbit IgG (H+L) A11036 |
| Astrocytes and neurons | Anti EAAT2 | mouse IgG2b | santa cruz | sc-365634 | dilution 1:500 | Alexa Fluor 647 Goat α-Mouse IgG (H+L) A21236 |
| Astrocytes and neurons | Anti Vimentin | chicken | abcam | ab24525 9822 | dilution 1:2200 | Goat α-chicken 568 abcam ab175477 |
| Astrocytes and neurons | Anti TUJ1 | mouse | Biolgened | 801201 | dilution 1:500 | Alexa Fluor 647 Goat α-Mouse IgG (H+L) A21236 |
| Astrocytes and neurons | Anti MAP2 | mouse | sigma | M4403 2ML | dilution 1:1000 | Alexa Fluor 647 Goat α-Mouse IgG (H+L) A21236 |
| Astrocytes and neurons | Anti GFAP | rabbit | Millipore | AB5804 | dilution 1:500 | Alexa Fluor 568 Goat α-Rabbit IgG (H+L) A11036 |
| Astrocytes and neurons | Anti TH | rabbit | santa cruz | sc-14007 | dilution 1:50 | Alexa Fluor 568 Goat α-Rabbit IgG (H+L) A11036 |
| Astrocytes and neurons | Anti HNF-3β (RY-7) [FoxA2] | mouse | santa cruz | sc-101060 | dilution 1:50 | Alexa Fluor 647 Goat α-Mouse IgG (H+L) A21236 |
| Astrocytes and neurons | S100β | rabbit | abcam | ab868 | dilution 1:50 | Alexa Fluor 568 Goat α-Rabbit IgG (H+L) A11036 |
| Astrocytes and neurons | Lamp1 | mouse | abcam | ab2296838 | dilution 1:1000 | Alexa Fluor 647 Goat α-Mouse IgG (H+L) A21236 |
| Astrocytes and neurons | Tom20 (FL-145) | rabbit | santa cruz | sc-11415 | dilution 1:1000 | Alexa Fluor 568 Goat α-Rabbit IgG (H+L) A11036 |
| Microglia | Iba1 | rabbit | FUJIFILM | 019-19741 | dilution 1:500 | Alexa Fluor 647 goat α-Rabbit IgG A27040 |
| T cells | FITC Mouse Anti-Human CD3 (clon UCHT1) | mouse | BD Pharmingen | 581806 | dilution 1:100 | |
| T cells | APC Mouse Anti-Human CD4 (clon L200) | mouse | BD Pharmingen | 551980 | dilution 1:100 | |

| | | | | | |
|---|---|---|---|---|---|
| T cells | PE Mouse Anti-Human CD8 (Clon RPA-T8) | mouse | BD Pharmingen | 561949 | dilution 1:100 |
| Astrocytes, T cells | DJ-1 (D29E5) | rabbit | cell signaling | mAb#5933 | dilution 1:1500 |
| Astrocytes, T cells | Beta actin | mouse | cell signaling | mAb#3700 | dilution 1:20000 |
| Human brain tissue | GFAP | | Sigma-HPA063513 | | dilution 1:800 |
| Human brain tissue | Aldoc | | Sigma-HPA003282 | | dilution 1:1000 |
| Human brain tissue | Iba1 | | Wako-019-19741 | | dilution 1:1000 |
| Biotin Switch Assay | DLAT Polyclonal Antibody | rabbit | Invitrogen | # PA5-59298 | dilution 1:250 |
| Biotin Switch Assay | DLST Polyclonal Antibody | rabbit | Invitrogen | # PA5-51794 | dilution 1:250 |
| Biotin Switch Assay | Vinculin (E1E9V) monoclonal anyobody | rabbit | cell signaling | mAb#13901 | dilution 1:750 |

### Suppl. Figure 1

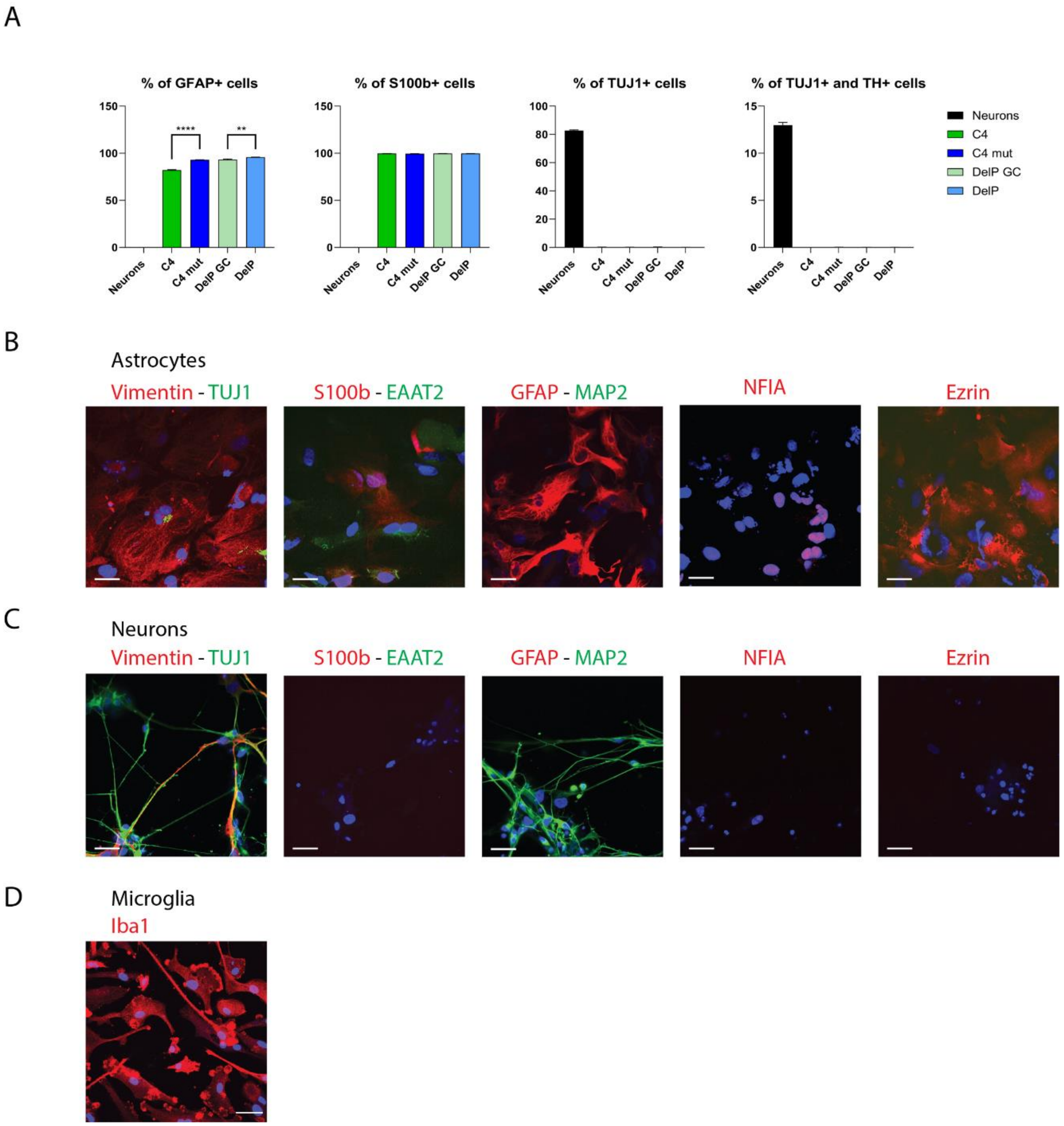

**Supplementary Figure 1: A:** Characterization of astrocytes by FACS. Almost all cells were GFAP and S100b positive. No neuronal contamination of astrocytic culture as assessed by TUJ1 staining. N=3-5. Error bars show SEM. Two-tailed paired T test was used. p <0.0001 = ****, p <0.001 = ***, p <0.01 = **, p <0.05 = *. **B and C:** Characterization of astrocytes by ICC. The cells showed an astrocytic morphology, and the majority stained positive for canonical astrocyte markers like GFAP, S100b, Vimentin, EAAT2, NFIA, ID3, EZRIN and no neuronal contamination as assessed by markers for TUJ1 and MAP2. **D:** Characterization of microglia by ICC. The cells showed microglia morphology and stained positive for the microglia marker Iba1.

**Suppl. Figure 2**

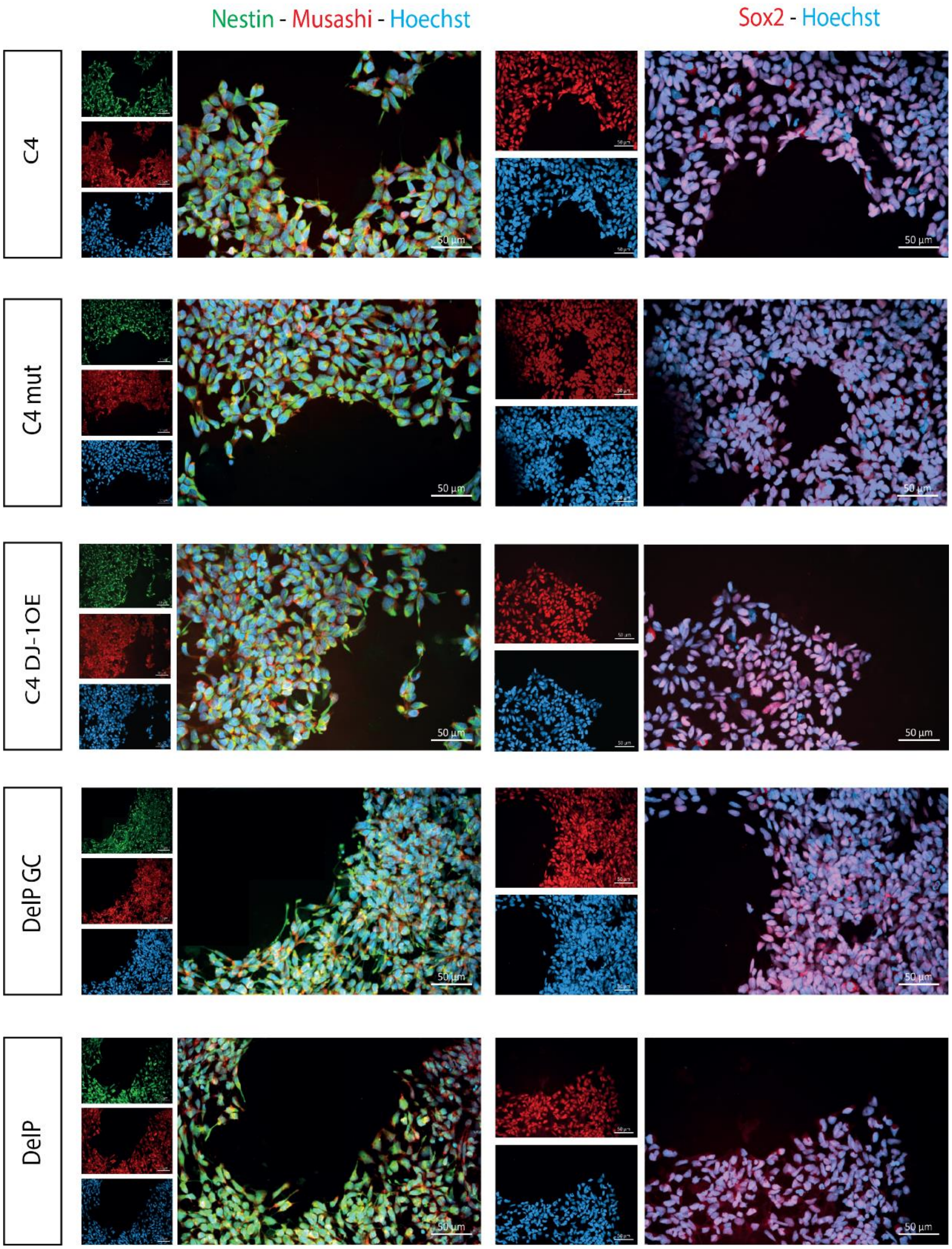

**Supplementary Figure 2:** Characterization of smNPCs by ICC.

**Suppl. Figure 3**

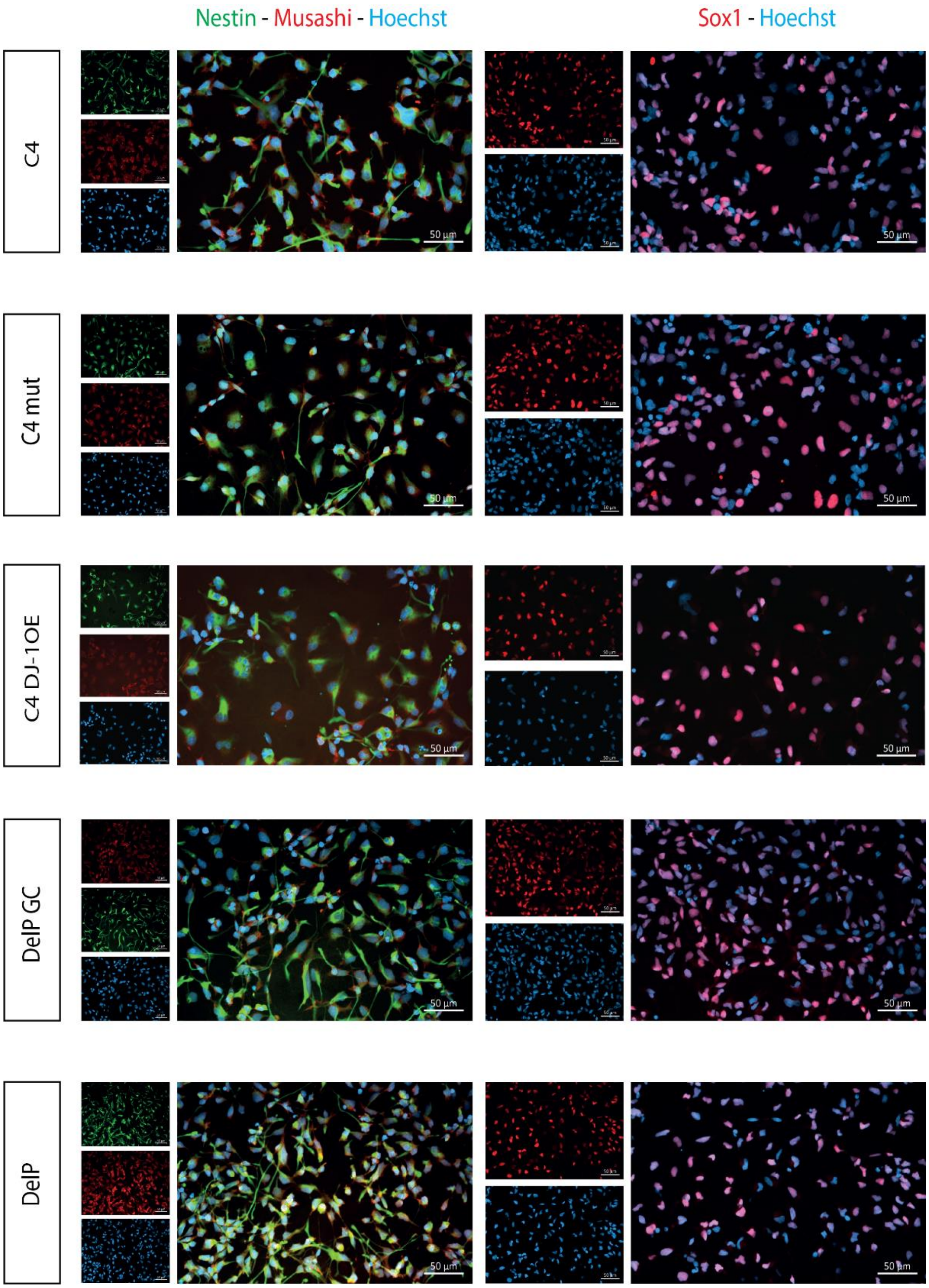

**Supplementary Figure 3:** Characterization of hNSCs by ICC.

**Suppl. Figure 4**

A

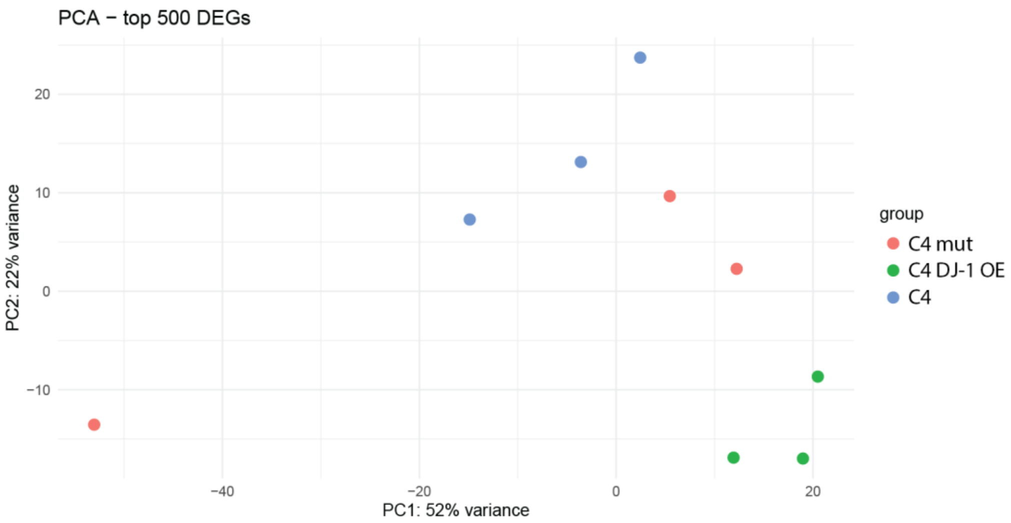

**Supplementary Figure 4:** RNAseq analysis of astrocytes. **A:** Principal Component Analysis (PCA) plot for RNA-seq data in each genotype based on top 500 DEGs. Three biological replicates for each cell line are represented separately.

### Suppl. Figure 5

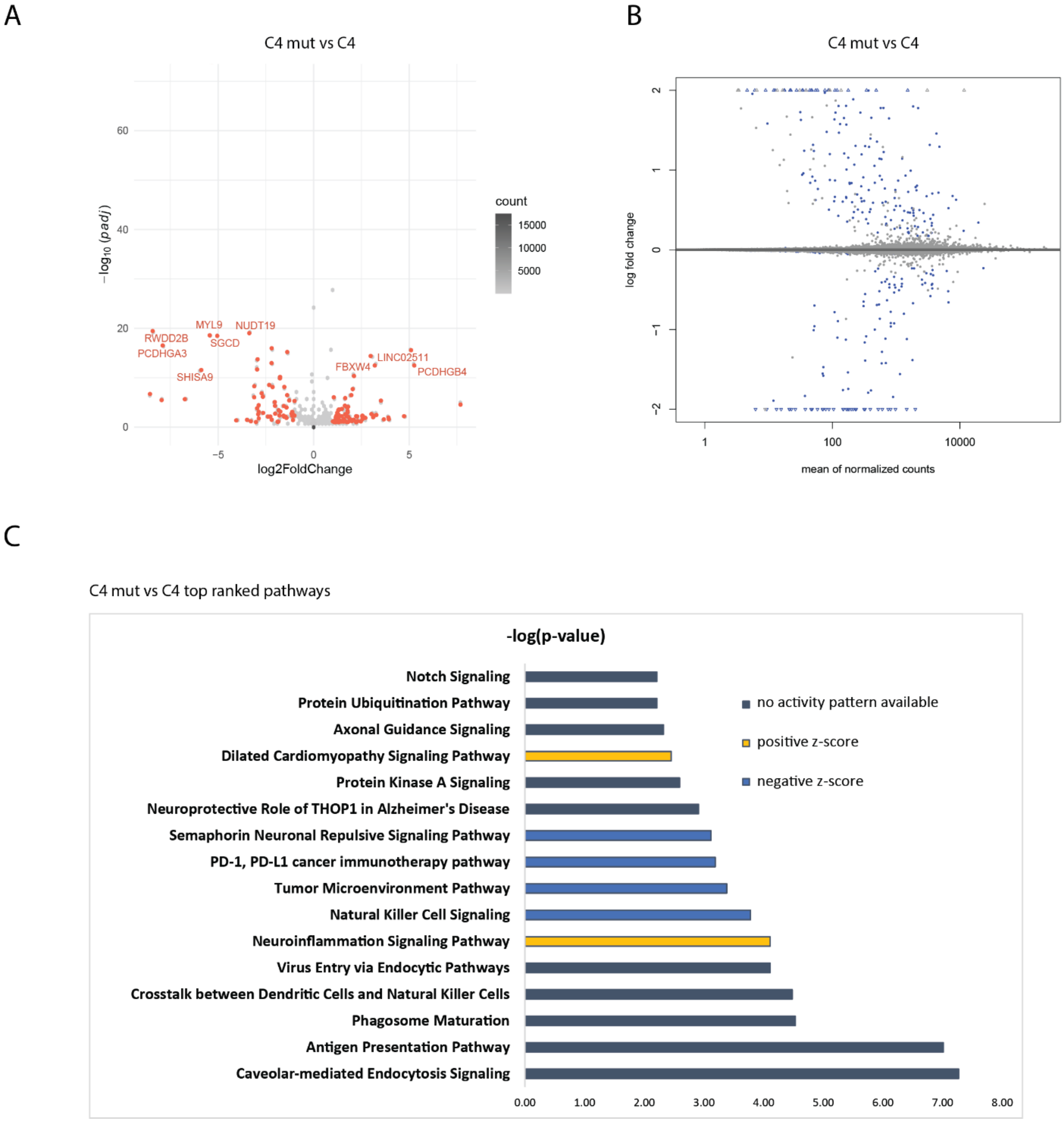

**Supplementary Figure 5:** RNAseq analysis of C4 mut astrocytes and C4 wildtype astrocytes. **A:** Volcano plot. Red dots highlight DEG with log2 fold change of +- - and padj < 0.05. gene names are reported for -log10(adjusted-pval) > 10 and abs(logFC) >= 3. **B:** MA-plot obtained after schrinkage of logFC by apeglm, blue dots highlight DEGs with padj < 0.05. **C:** Top ranked pathways of IPA analysis showing only −log(p-value) > 2 revealed that different pathways were enriched and that especially neuroinflammation pathways were activated (z-score >0.5).

**Suppl. Figure 6**

A

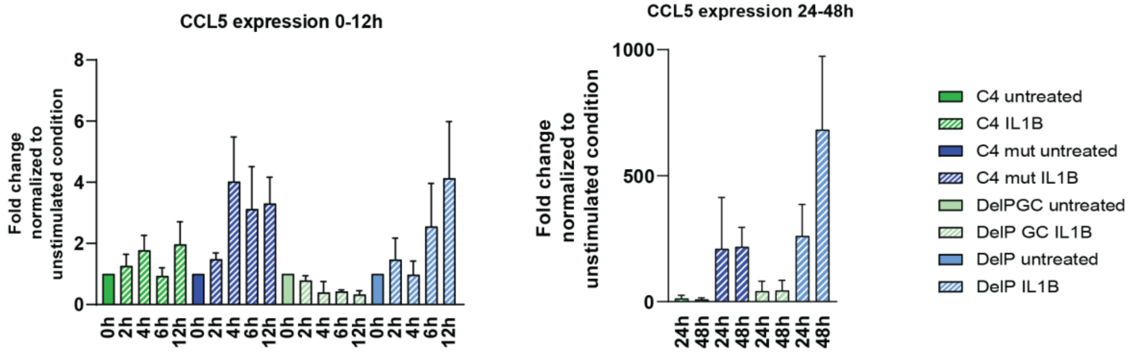

B

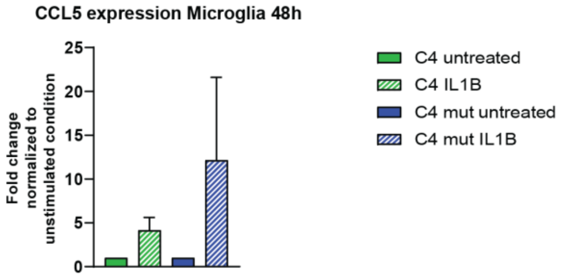

**Supplementary Figure 6:** CCL5 mRNA expression upon IL-1β treatment (10 ng/ml) at indicated time points. **A:** Expression levels of CCL5 mRNA in astrocytes at 0h, 2h, 4h, 6h, 12h, 24h, and 48h after IL-1β stimulation assessed by qPCR. N= 3. **B:** Expression levels of CCL5 mRNA in iPSC-derived microglia 48h after IL-1β stimulation assessed by qPCR. N= 3.

**Suppl. Figure 7**

A

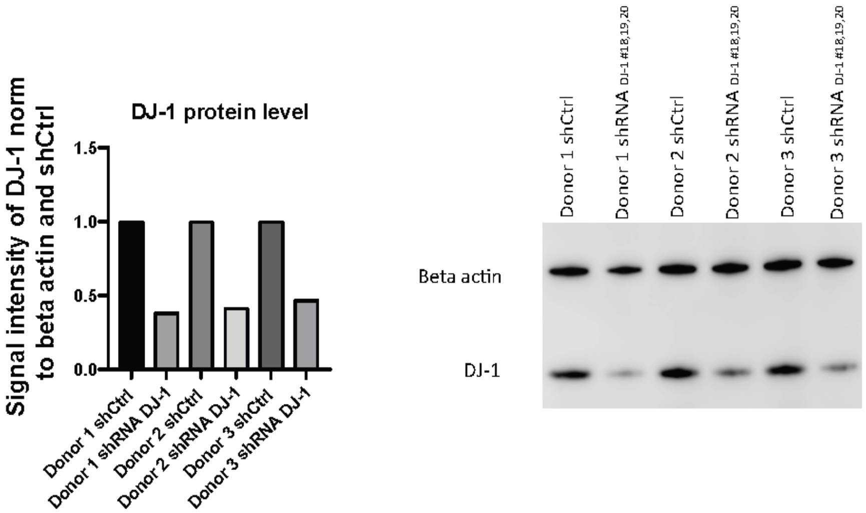

B

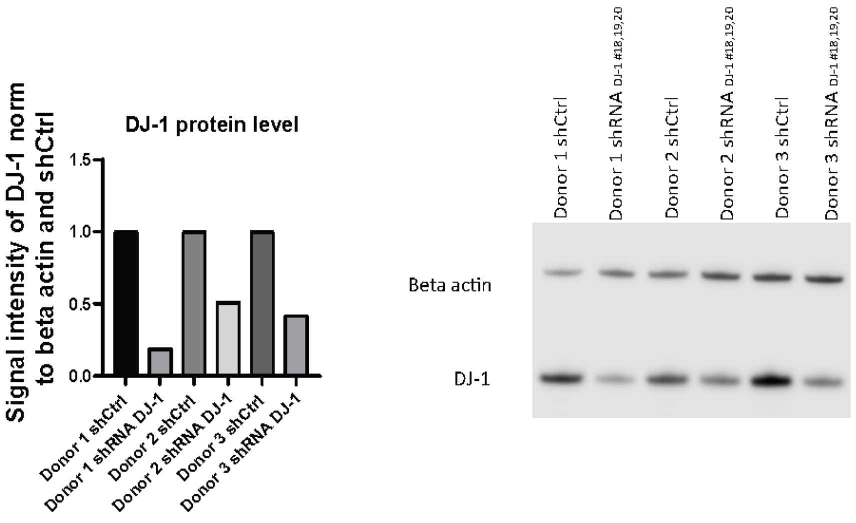

**Supplementary Figure 7: A+B:** DJ-1 knockdown in T cells prior to T cell migration assay using shRNAs #18 and #20 together. Each PBMC donor represents one independent biological replicate that was used for the respective different astrocyte differentiations. N=3. Panel A shows the knockdown for the T cell migration assay performed with C4 and C4 mut astrocytes, panel B for the assay with DelP GC and DelP astrocytes.

**Suppl. Figure 8**

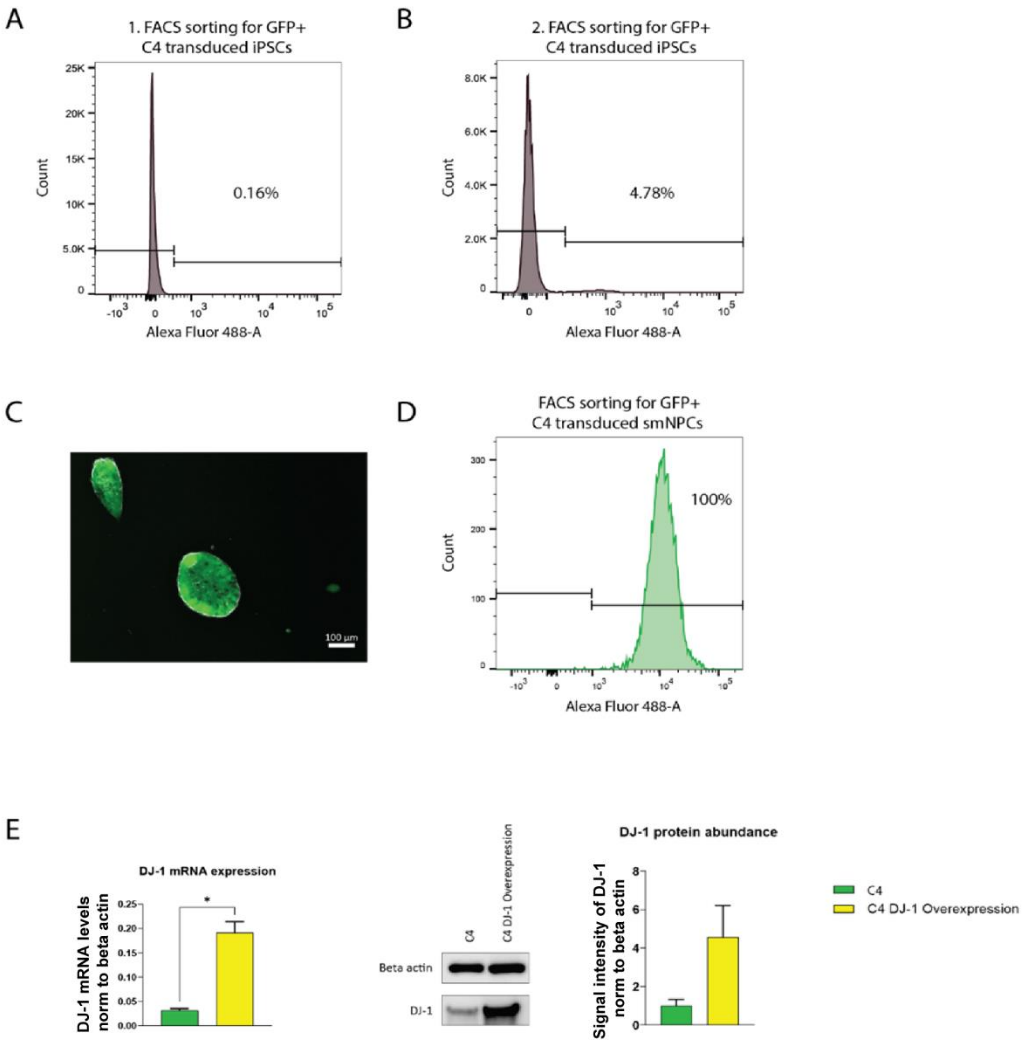

**Supplementary Figure 8: A:** FACS sorting for GFP+ cells of transduced C4 wildtype iPSC with GFP containing lentiviral construct for DJ-1 overexpression. For the first sort, 0.16% of the cells were GFP+. **B:** Second sort of transduced iPSC resulted in 4.78% GFP+ cells. **C:** GFP+ iPSC colonies after the second sort. **D:** GFP+ iPSCs were differentiated into smNPCs and sorted for GFP+ cells again which resulted in 100% GFP+ cells. **E:** Characterization of DJ-1 overexpression smNPCs showed increased DJ-1 mRNA and protein level.

**Suppl. Figure 9**

A

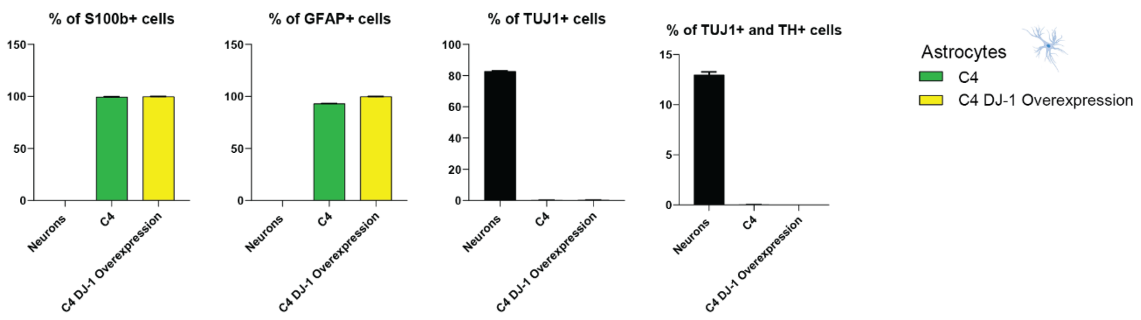

**Supplementary Figure 9: A:** FACS-based characterization of astrocytes showed that astrocytes express astrocyte markers GFAP and S100b, but no neuronal contamination due to absence of TUJ1 staining.

### Suppl. Figure 10

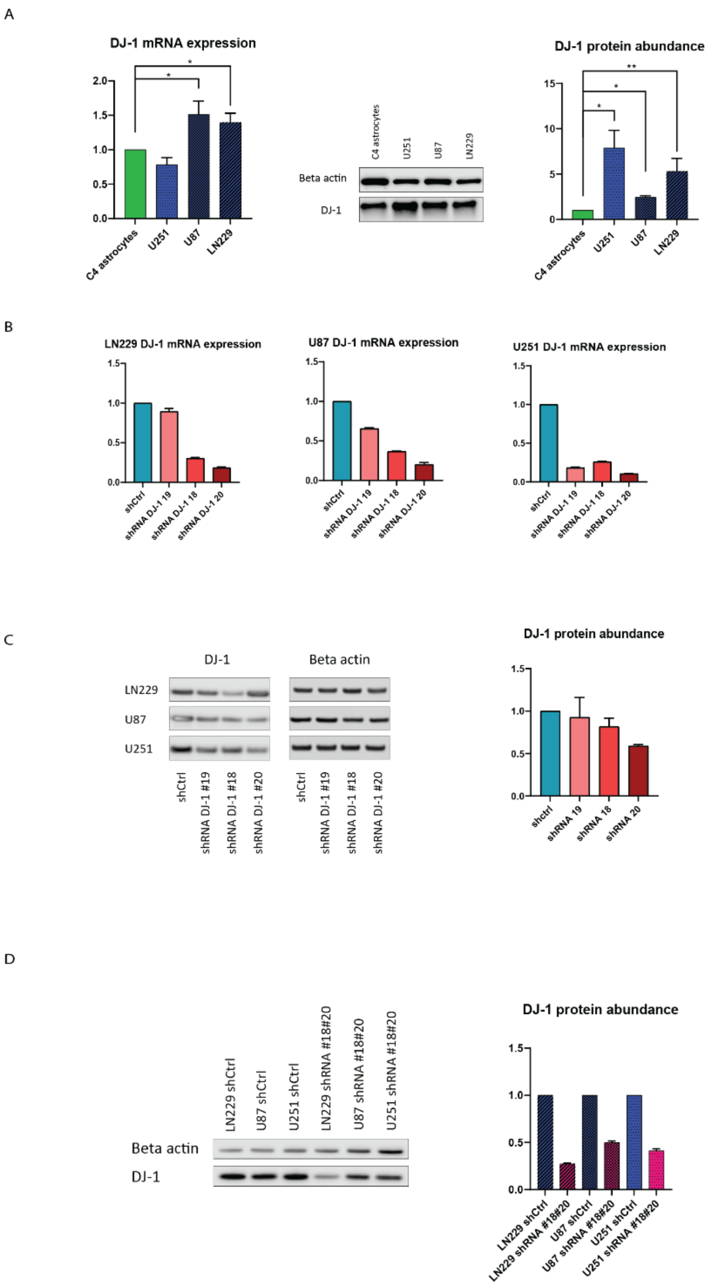

**Supplementary Figure 10: A:** GBM cell lines have upregulated DJ-1 mRNA and protein levels when compared to C4 astrocytes. **B:** shRNA mediated knockdown of DJ-1 reduced DJ-1 mRNA level. shRNA #20 had the strongest knockdown effect in all the lines and reduced the DJ-1 mRNA levels to around 20% compared to shCtrl. **C:** DJ-1 protein levels upon knockdown reduced to 50-60% compared to shCtrl. **D:** DJ-1 protein levels upon stable knockdown reduced to 30-50% compared to shCtrl.

**Suppl. Figure 11**

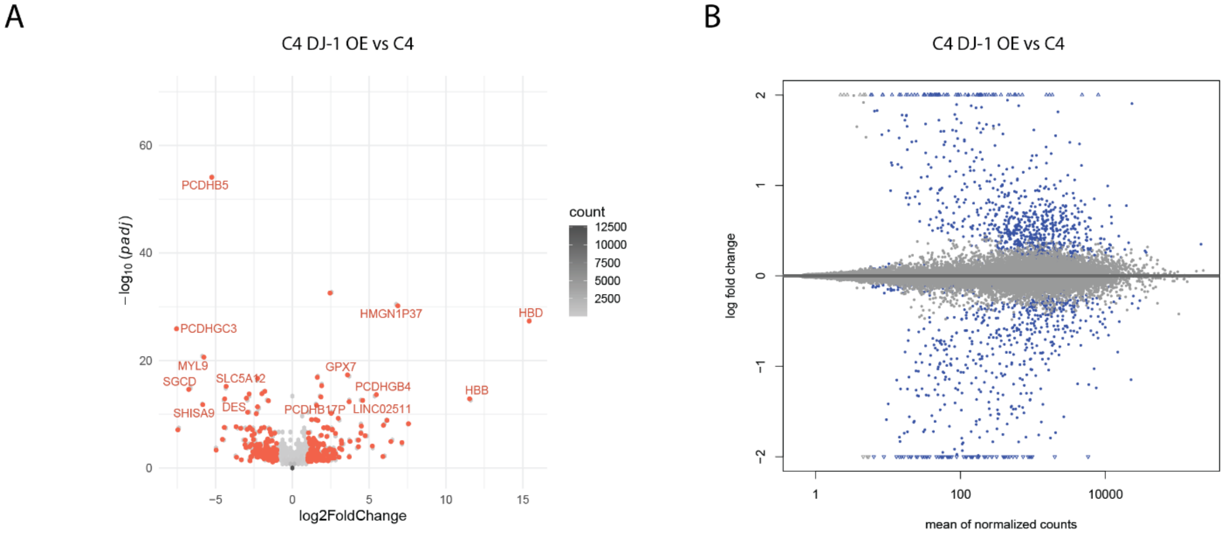

**Supplementary Figure 11:** RNAseq analysis of C4 DJ-1 OE astrocytes and C4 wildtype astrocytes. **A:** Volcano plot. Red dots highlight DEG with log2 fold change of +- - and padj < 0.05. gene names are reported for -log10(adjusted-pval) > 10 and abs(logFC) >= 3. **B:** MA-plot obtained after schrinkage of logFC by apeglm, blue dots highlight DEGs with padj < 0.05.

**Suppl. Figure 12**

A

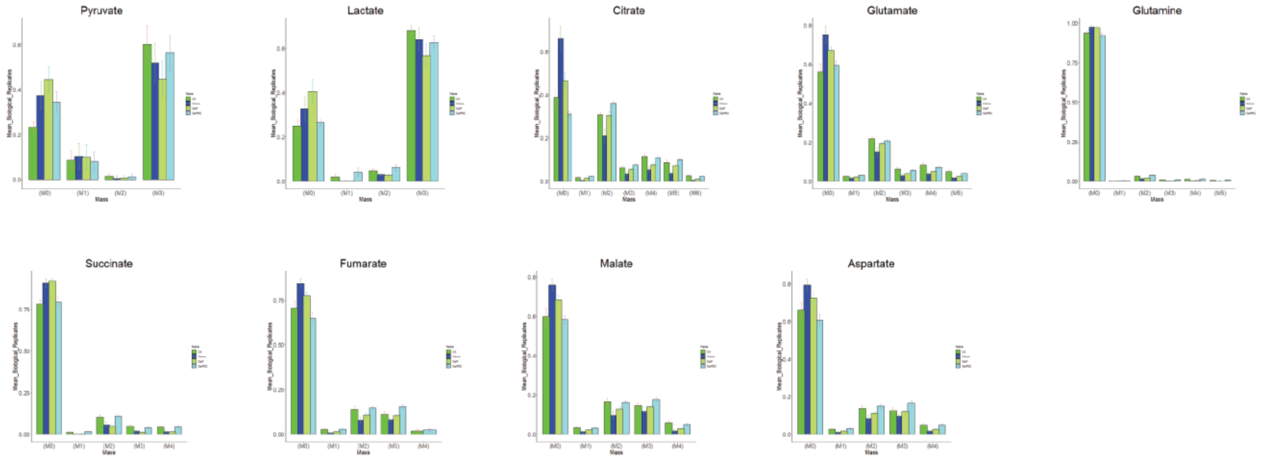

B

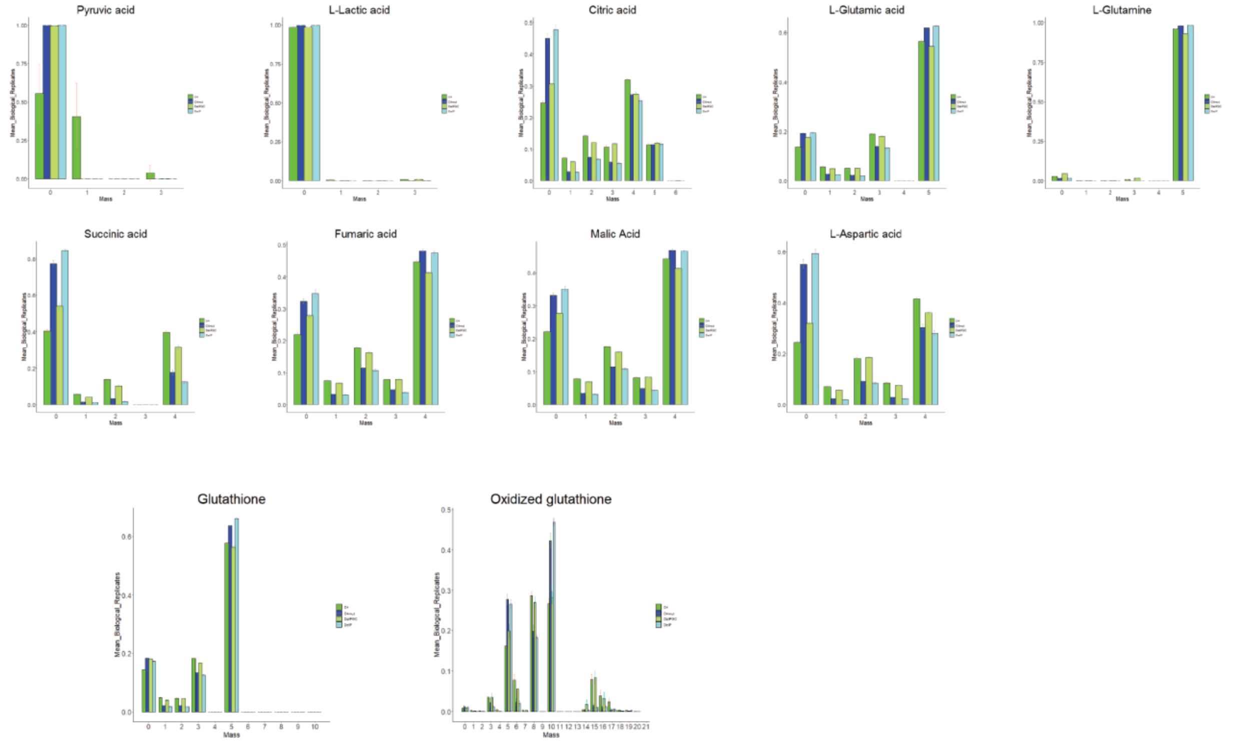

**Supplementary Figure 12: Full MIDs for experiments presented in Figure 3. A:** Analysis of glucose metabolism using [U-13C6]Glucose tracing. N=3. 13C incorporation in metabolites was analyzed by GC-MS, resulting in heavier metabolites (M1+x), whereas no $^{13}C$ incorporation corresponds to M0. **B:** [U-13C5]Glutamine tracing results. N=3.

**Suppl. Figure 13**

A

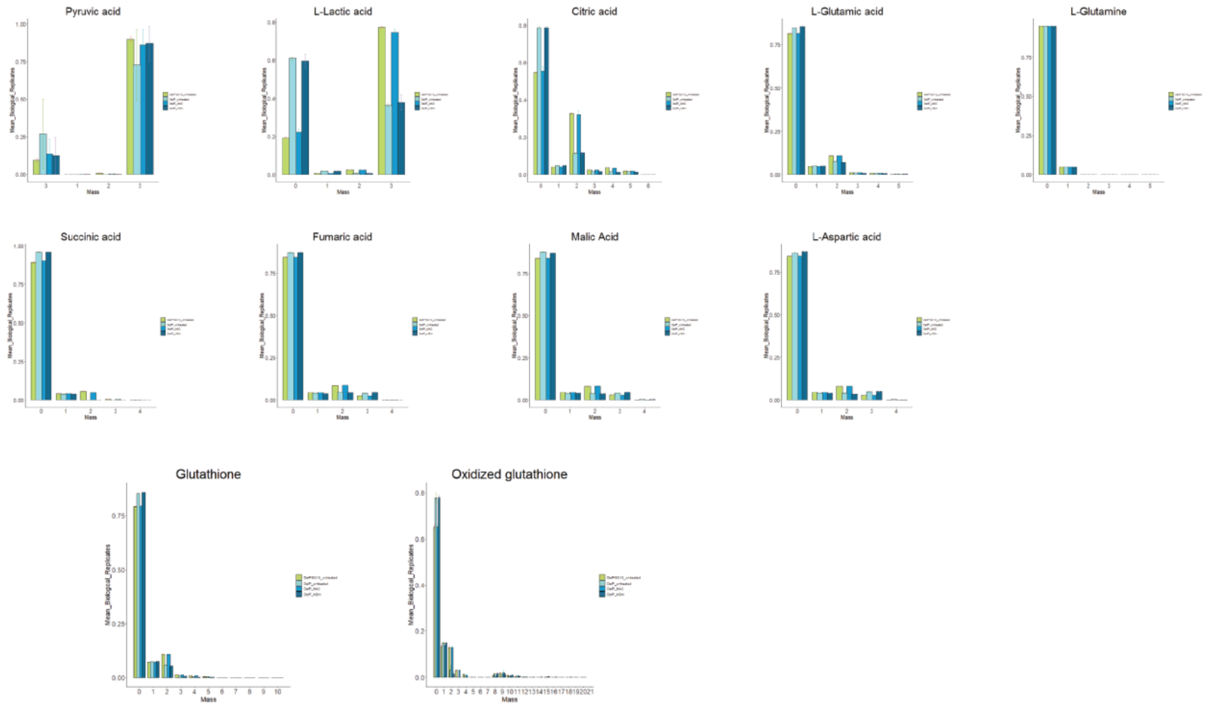

B

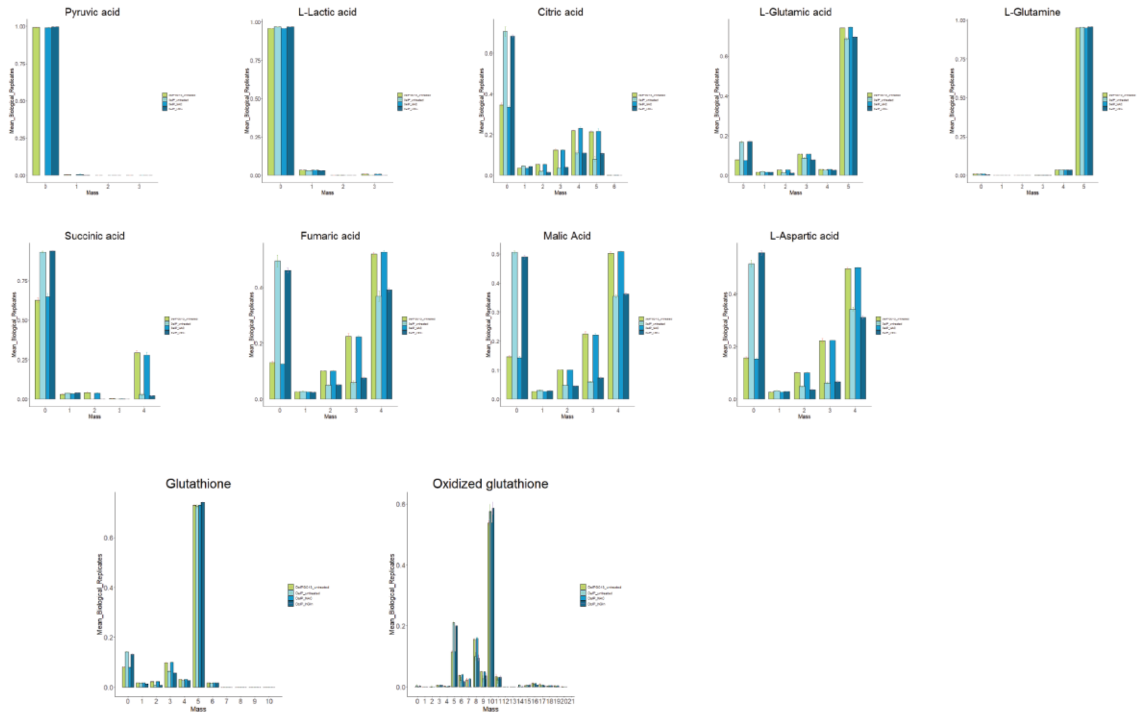

**Supplementary Figure 13: Full MIDs for experiments presented in Figure 7. A:** Analysis of glucose metabolism using [U-13C6]Glucose tracing. N=3. 13C incorporation in metabolites was analyzed by GC-MS, resulting in heavier metabolites (M1+x), whereas no $^{13}C$ incorporation corresponds to M0. **B:** [U-13C5]Glutamine tracing results. N=3.

**Suppl. Figure 14**

A

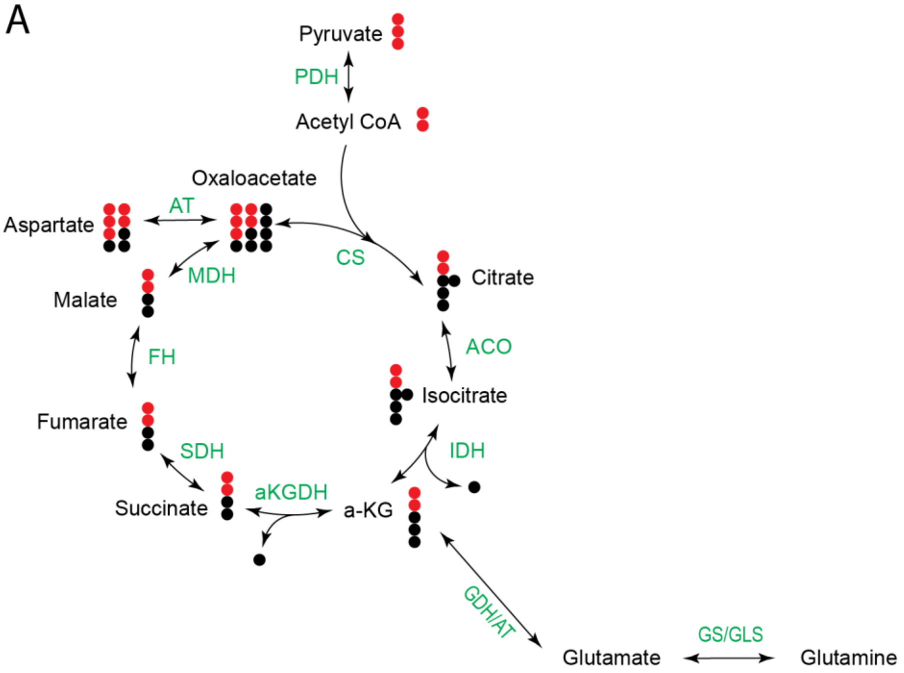

**Supplementary Figure 14: A:** Principle of [U-13C6]Glucose tracing. 13C (red) incorporation in metabolites is analyzed by GC-MS, resulting in heavier metabolites (M1+x), whereas no 13C incorporation corresponds to M0.

**Suppl. File 1**

Whole scans of immunohistochemically stained brain slides. Files can be viewed using QuPath.
**A:** Control brain slide stained for Aldoc
**B:** Control brain slide stained for GFAP
**C:** Control brain slide stained for Iba1
**D:** DJ-1 patient brain slide stained for Aldoc
**E:** DJ-1 patient brain slide stained for GFAP
**F:** DJ-1 patient brain slide stained for Iba1
